# Toxin-antitoxin gene pairs found in Tn*3* family transposons appear to be an integral part of the transposition module

**DOI:** 10.1101/848275

**Authors:** Gipsi Lima-Mendez, Danillo Oliveira Alvarenga, Karen Ross, Bernard Hallet, Laurence Van Melderen, Alessandro M. Varani, Michael Chandler

## Abstract

Much of the diversity of prokaryotic genomes is contributed by the tightly controlled recombination activity of transposons (Tn). The Tn*3* family is arguably one of the most widespread transposon families. Members carry a large range of passenger genes incorporated into their structures. Family members undergo replicative transposition using a DDE transposase to generate a cointegrate structure which is then resolved by site-specific recombination between specific DNA sequences (*res*) on each of the two Tn copies in the cointegrate. These sites also carry promoters controlling expression of the recombinase and transposase. We report here that a number of Tn*3* members encode a type II toxin-antitoxin (TA) system, typically composed of a stable toxin and a labile antitoxin that binds the toxin and inhibits its lethal activity. This system serves to improve plasmid maintenance in a bacterial population and, until recently, was believed to be associated with bacterial persistence. At least six different TA gene pairs are associated with various Tn*3* members. Our data suggest that several independent acquisition events have occurred. In contrast to most Tn*3* family passenger genes which are generally located away from the transposition module, the TA gene pairs abut the *res* site upstream of the resolvase genes. Although their role when part of Tn*3* family transposons is unclear, this finding suggests a potential role for the embedded TA in stabilizing the associated transposon with the possibility that TA expression is coupled to expression of transposase and resolvase during the transposition process itself.

**Importance:** Transposable Elements (TEs) are important in genetic diversification due to their recombination properties and their ability to promote horizontal gene transfer. Over the last decades, much effort has been made to understand TE transposition mechanisms and their impact on prokaryotic genomes. For example, the Tn*3* family is ubiquitous in bacteria, moulding their host genomes by the *paste-and-copy* mechanism. In addition to the transposition module, Tn*3* members often carry additional passenger genes (e.g., conferring antibiotic or heavy metal resistance and virulence) and three were previously known to carry a toxin-antitoxin (TA) system often associated with plasmid maintenance; however, the role of TA systems within the Tn*3* family is unknown. The genetic context of TA systems in Tn*3* members suggests that they may play a regulatory role in ensuring stable invasion of these Tn during transposition.

## Introduction

Members of the Tn*3* transposon family form a tightly knit group having related transposase genes and related DNA sequences at their ends. However, they are highly diverse in the range of passenger genes they carry (see (1)) (Figure 1). The basic Tn*3-*family transposition module is composed of transposase and resolvase genes and two ends with related terminal inverted repeat DNA sequences, the IRs, of 38-40 bp or sometimes even longer (2). They encode a large (∼1000 aa) DDE transposase, TnpA, significantly longer than the DDE transposases normally associated with Insertion Sequences (IS) (see (3)). The TnpA transposase catalyzes DNA cleavage and strand transfer reactions necessary for formation of a cointegrate transposition intermediate during replicative transposition (4). The cointegrate is composed of donor (with the transposon) and target (without the transposon) circular DNA molecules fused into a single circular molecule and separated by two directly repeated transposon copies, one at each donor-target junction (4). Phylogenetic analysis based on TnpA sequence identified 7 clusters or sub-groups named after representative transposons: Tn*3*, Tn*21*, Tn*163*, IS*1071*, IS*3000*, Tn*4430* and Tn*4651* (1, 5). A second feature of members of this transposon family is that they carry short internal (∼100-150 bp) DNA segments, at which site-specific recombination between each of the two Tn copies occurs to “resolve” the cointegrate into individual copies of the transposon donor and the target molecules each containing a single transposon copy (1).

**Figure 1:**
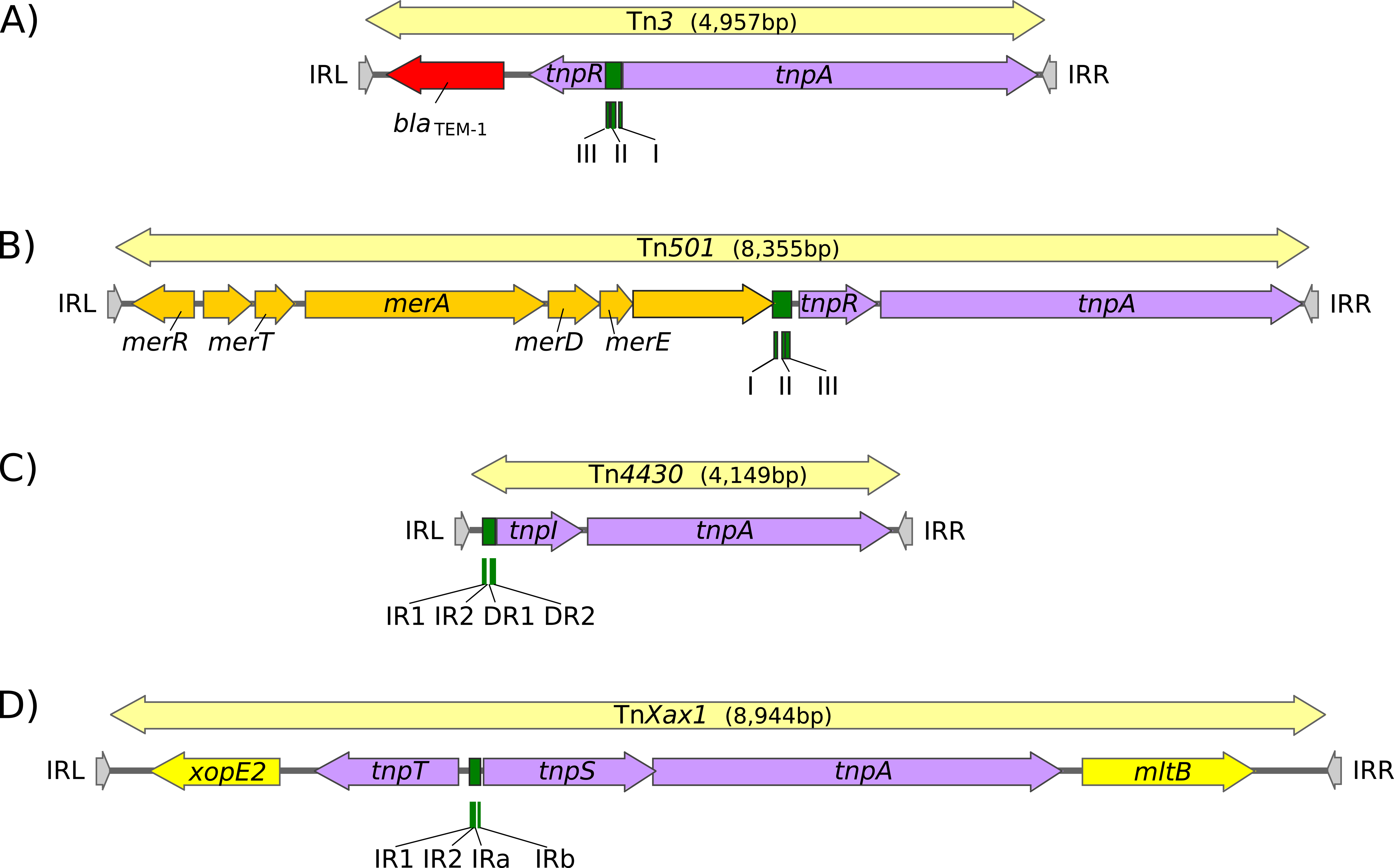
Tn*3*-family architectures. This figure illustrates the major different types of gene arrangement in members of the Tn*3* family. Transposons are shown as pale yellow boxes ending in arrowheads. The transposon length in base pairs is indicated. Terminal inverted repeats, IR, are indicated by grey arrowheads (IRL and IRR respectively, labelled by convention with respect to the direction of *tnpA* transcription from left to right). Recombination sites (*res, irs, rst*) are shown in green, transposition genes in purple, and passenger genes in red (antibiotic resistance genes), chrome yellow (heavy metal resistance genes) and strong yellow (plant pathogenicity genes). A) Tn*3*: Accession # V00613 (Tn*3* sub-group). Carries the *bla_TEM-1a_* beta-lactamase gene and divergent serine recombinase/resolvase, *tnpR*, and transposase, *tnpA*, genes. The recombination site, *res*, composed of three sub sequences I, II and III, is located between *tnpR* and *tnpA*. with site III proximal to *tnpR*. Recombination occurs within site I. B) Tn*501*: Accession # Z00027 (Tn*21* sub-group). Carries an operon containing mercury resistance genes (*mer*) and collinear serine recombinase/resolvase, *tnpR*, and transposase, *tnpA*, genes. The *res* site is located upstream of *tnpR*. It has a similar organization as that of Tn*3* with site III proximal to *tnpR*. Recombination occurs within site I. C) Tn*4430*: Accession # X07651.1 (Tn*4430* sub-group). Carries no known passenger genes. A tyrosine recombinase/resolvase, *tnpI*, and transposase, *tnpA*, genes are colinear and the recombination site, *irs*, is located upstream of and proximal to the resolvase gene with four subsites: inverted repeats IR1 and IR2 and direct repeats DR1 and DR2. Recombination occurs at the recombination core site IR1-IR2. D) Tn*Xax1*: Accession #AE008925 (Tn*4651* sub-group). Carries two passenger genes involved in plant pathogenicity located at the left (*xopE*) and right (*mlt*) ends of the transposon. The resolvase has two components: a tyrosine recombinase, *tnpT* and a helper protein, *tnpS* expressed divergently. The res site, *rst*, is located between *tnpT* and *tnpS* and is composed of two pairs of inverted repeats IR1 and IR2 and IRa and IRb. Recombination occurs at the IR1-IR2 inverted repeat.

This highly efficient recombination system is assured by a transposon-specified site-specific recombinase: the resolvase. There are at present three known major resolvase types: TnpR, TnpI and TnpS+TnpT (Figure 1), distinguished, among other features, by the catalytic nucleophile involved in DNA phosphate bond cleavage and rejoining during recombination. TnpR is a classic serine (S)-site-specific recombinase (e.g., (6)); TnpI is a tyrosine (Y) recombinase (7)(see (1)) and TnpS+TnpT is a heteromeric resolvase combining a tyrosine recombinase, TnpS, and a divergently expressed helper protein, TnpT, with no apparent homology to other proteins (8, 9). The *tnpR* gene can be either in the same orientation or opposite orientation as *tnpA*. In the former case the *res* site lies upstream of *tnpR* and in the latter case, between the divergent *tnpR* and *tnpA* genes. For relatives encoding TnpS and TnpT, the corresponding genes are divergent and the *res* (*rst*) site lies between *tnpS* and *tnpT*. Examples of these architectures are shown in Figure 1. Each *res* includes a number of short DNA sub-sequences which are recognized and bound by the cognate resolvases. These are different for different resolvase systems and called *res* (for *res*olution site) or IRS (10), *irs* (for internal recombination site, (11)) or *rst* (for *<u>r</u>*esolution site tnp*<u>S</u>* tnp*<u>T</u>*; (8))(see below)*. res* sites that have been analyzed also include promoters that drive both transposase and resolvase expression (see (1, 10, 12)). Indeed, TnpR from Tn*3* itself was originally named for its ability to repress transposase expression by binding to these sites (13, 14).

The diversity of these Tn resides in the variety of other mobile elements that have been incorporated into their structures such as IS and integrons, as well as other Tn*3* family members (see (1)) and of their passenger genes. The most notorious of these passenger genes are those for antibiotic and heavy metal resistance although other genes involved in virulence functions for both animals and plants (e.g., Figure 1) or in organic catabolite degradation also form part of the Tn*3* family passenger gene arsenal.

While studying *Xanthomonas citri*, a principal pathogen of citrus trees and an important economic problem (e.g. (15)), we had identified a number of Tn*3*-family structures in pXac64, a conjugative plasmid carrying a variety of pathogenicity and virulence genes (Figure 2) (2, 16). An interesting observation was that one of the Tn*3*-related transposons, Tn*Xc4*, carries a toxin-antitoxin (TA) system belonging to the type II TA class (17).

**Figure 2.**
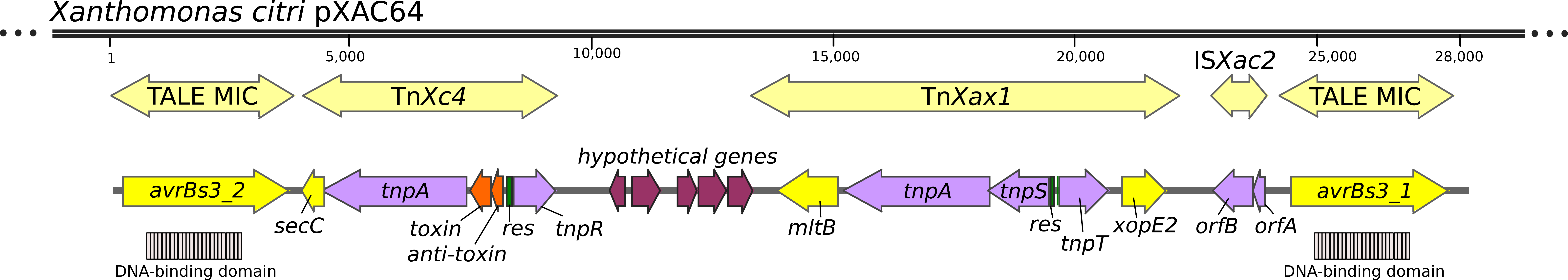
An annotated map of plasmid pXAC64 (accession # CP004400) from *Xanthomonas citri*. The figure shows a section of the plasmid carrying four Tn*3* derivative transposons and one Insertion Sequence, IS*Xac2*. Derivatives located to the left and right are MICs (Minimal Insertion Cassettes; (56)), which are devoid of transposition genes. These include TALE (Transcription Activator Like Elements) (yellow) responsible for pathogenicity, which, in turn, include an array of peptide repeats (DNA-binding domain in grey). Between the flanking MICs are two complete Tn*3*-family transposons. Tn*Xax1* carries a tnpS/T resolvase with an intervening *res* (*rst*) recombination site (green) and two genes (yellow) involved in plant-pathogen interaction. Tn*Xc4* includes a toxin antitoxin gene pair, in orange, and divergent *tnpA* and *tnpR* genes with an intervening *res* site (green). Coordinates in base pairs are shown on the external line.

Type II TA systems are generally composed of 2 proteins: a stable toxin and a labile antitoxin that binds the toxin and inhibits its lethal activity (see (18)). The antitoxin includes a DNA binding domain involved in promoter binding and negative regulation of TA expression. They are involved in plasmid maintenance in growing bacterial populations by a mechanism known as post-segregational killing. Upon plasmid loss, degradation of the labile antitoxin liberates the toxin from the inactive complex, which in turn is free to interact with its target and cause cell death. Recently, the Eva Top laboratory (19) while studying plasmid maintenance, observed that a relatively unstable plasmid, pMS0506, could be stabilised by transposition of a 7.1-kb Tn*3*-related transposon, Tn*6231*, from the non-self-transmissible plasmid pR28 (20) indigenous to *Pseudomonas moraviensis*. Further analysis revealed that Tn*6231* (which is reported to be 99% identical to Tn*4662*; (19)) also carried a type II TA gene pair that presumably stabilized the target plasmid.

Although the presence of TA systems in Tn*3*-family transposons had been noted previously (21, 22)(see (1)), neither the function of these systems within the Tn*3* family nor their genetic context have been examined. These initial observations prompted us to investigate whether TA systems have been acquired by other Tn*3*-family members in a similar way and to examine their possible involvement in Tn behavior.

## Results

### Identification of TA gene pairs in Tn*3* family members

As a first step, we undertook a detailed annotation of available Tn*3* family members in the ISfinder database (23) and also those listed in Nicolas et al. (1). We also searched NCBI for previously annotated Tn*3* family members (March 2018) and made use of an in-house script which searches for *tnpA*, *tnpR* and TA genes located in proximity to each other (Tn*3*finder: https://tncentral.proteininformationresource.org/TnFinder.html Tn*3*+TA_finder: https://github.com/danillo-alvarenga/tn3-ta_finder) to search complete bacterial genomes in the Refseq database at NCBI. Of 190 Tn*3* family transposons for which relatively complete sequence data (transposase, resolvase and generally both IRs) were available, 39 carry TA systems (Figure 3, colored squares; Tables 1 and S1). A phylogenetic tree based on similarity between the *tnpA* gene products is shown in Figure 3. Note that, with minor exceptions, the entire Tn*3* library conforms closely to the previously defined Tn*3*-family sub-groups (1). The majority of Tn*3*-family members encode a TnpR resolvase (Figure 3, purple circles) although several members of the Tn*163* sub-group carry the TnpS+TnpT resolvase (Figure 3, pink circles). Only three derivatives, Tn*5401*, Tn*Bth4* and Tn*4430* encode the TnpI resolvase (Figure 3, salmon circles).

**Figure 3.**
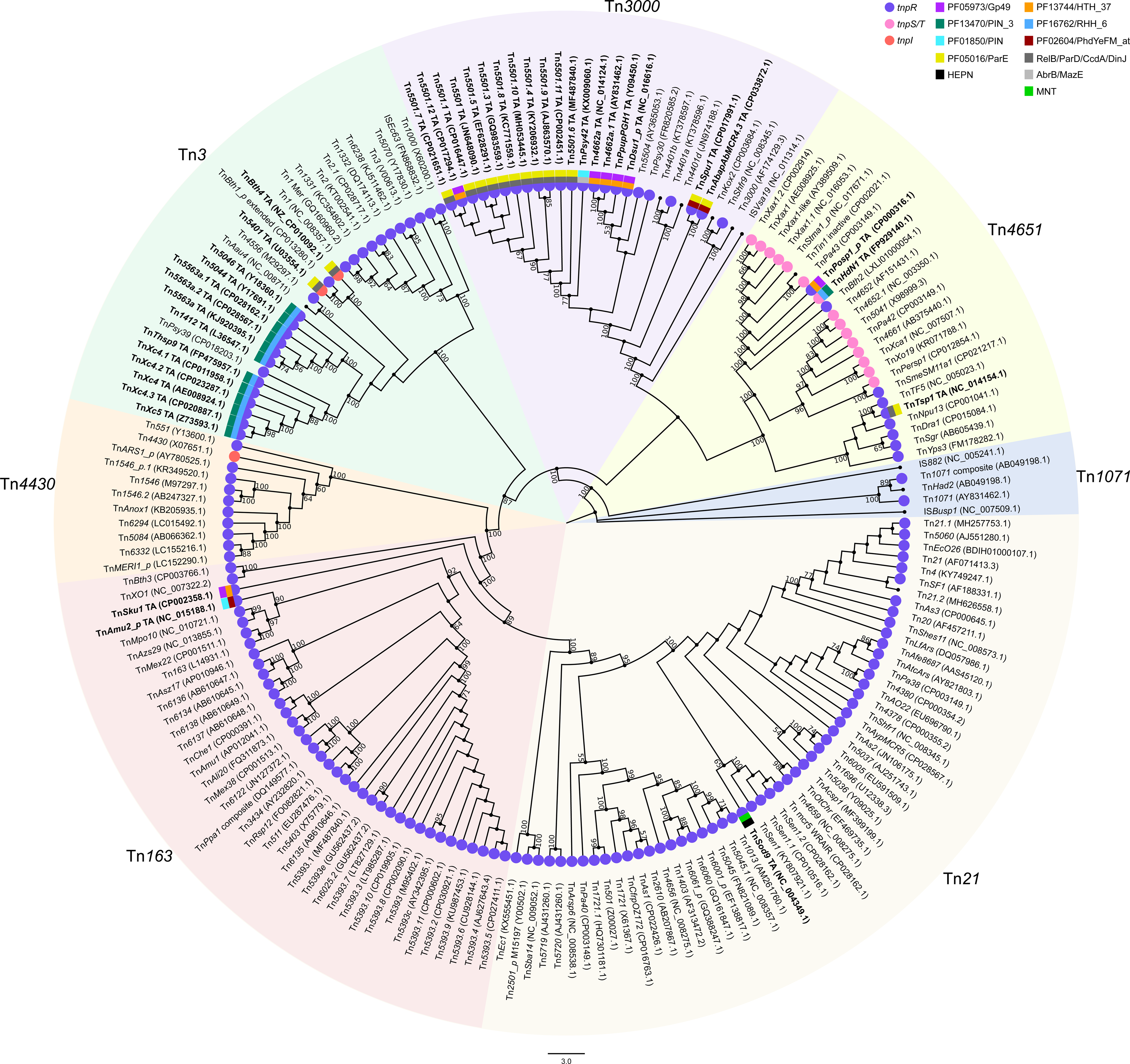
A phylogenetic tree of 190 Tn*3*-family members based on their TnpA sequences. We extracted Tn*3* family members from the ISfinder database which served to generate the sub-groups defined in Nicolas et al. (1). Many others were drawn from the literature and have been given official names (Tn followed by digits e.g. Tn*1234*: https://transposon.lstmed.ac.uk/tn-registry) while others were identified using Tn*3*_finder software (TnCentral: https://tncentral.proteininformationresource.org/TnFinder.html) and given temporary names. Each is associated with its GenBank accession number; the GenBank file contains either the extracted transposon or the DNA sequence from which it was extracted (e.g., DNA fragment, plasmid or chromosome). Numbers above the lines of each clade indicate the maximum likelihood bootstrap values. The sub-groups adhere closely to those defined by Nicolas et al. with some minor variations resulting from the significantly larger Tn sample. The majority of members carry *tnpR*, serine resolvases (purple circles). Those that include *tnpI* or *tnpT*/*tnpS* are indicated by salmon and pink circles, respectively. The TA gene pairs are indicated by coloured squares. Note that Tn5501.5 carries a mutation which truncates its toxin gene leaving the antitoxin intact. The outer squares represent the toxin and the inner squares, the antitoxin. The five toxin types are: Gp49 (PF05973), purple; PIN_3 (PF13470), dark green; PIN (PF01850), bright blue; ParE (PF05016), yellow; and HEPN, black. The antitoxins are: HTH_37 (PF13744), orange; RHH_6 (PF16762), blue; PhdYeFM_at (PF02604), magenta; RelB/ParD/CcdA/DinJ, dark grey; AbrB/MazE, light grey; MNT, bright green. The corresponding Tn names and accession numbers are highlighted in bold type for clarity. Note that the branches have been extended for clarity.

**Table 1.**
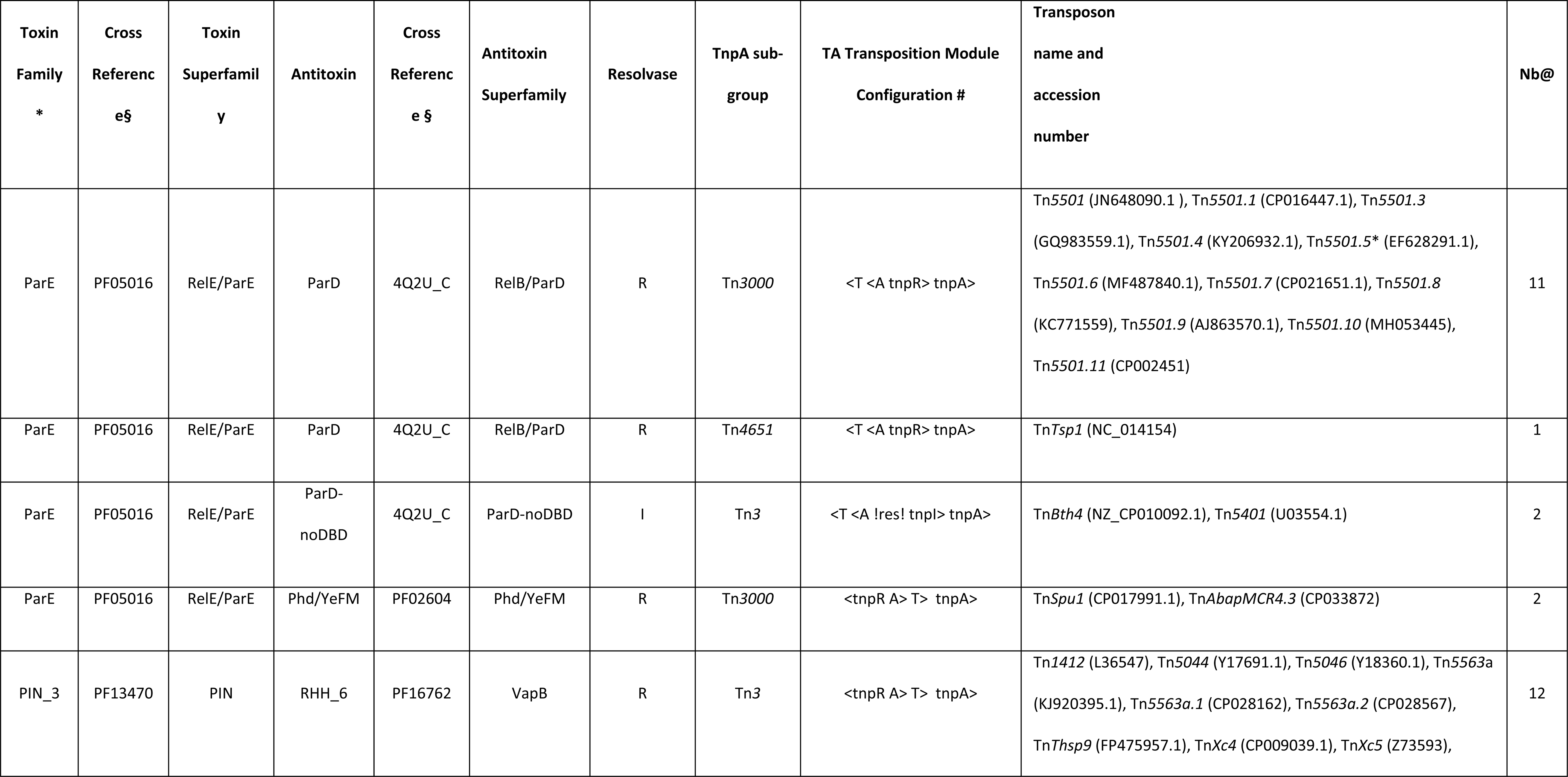
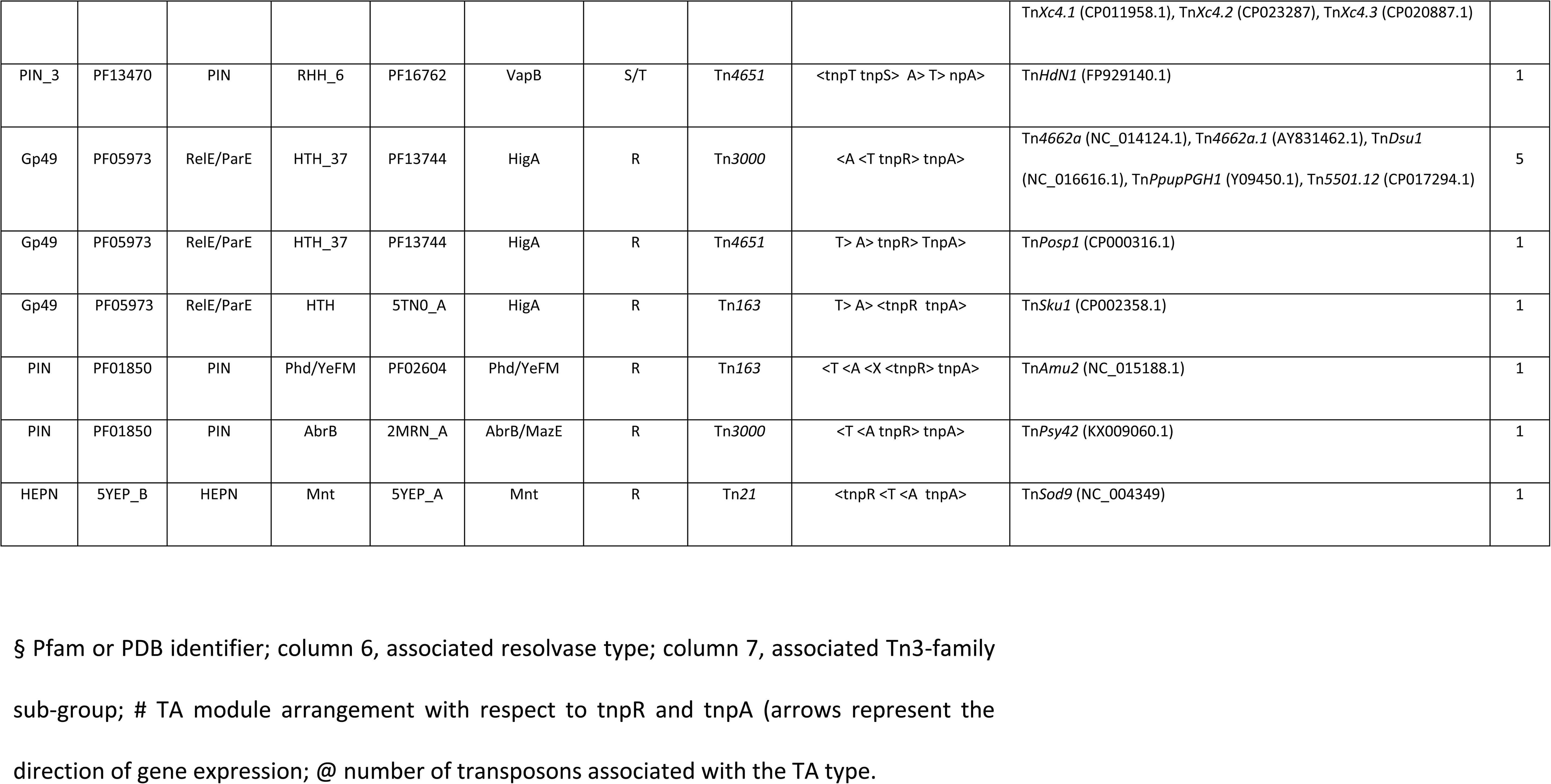
Tn*3* family TA systems and configuration of the TA operon with respect to the resolvase and TnpA genes.

The Tn carrying TA systems featured examples from all known combinations and orientation of transposase and resolvase genes (Figure 4). While most cases occurred in Tn*3* family members with *tnpR* resolvase genes, examples were also identified in transposons with *tnpS+tnpT* (Tn*Posp1_p* and Tn*HdN1.1*) and *tnpI* (Tn*5401* and Tn*Bth4*) genes (Figure 3). Illustrative examples are shown in Figure 4.

**Figure 4.**
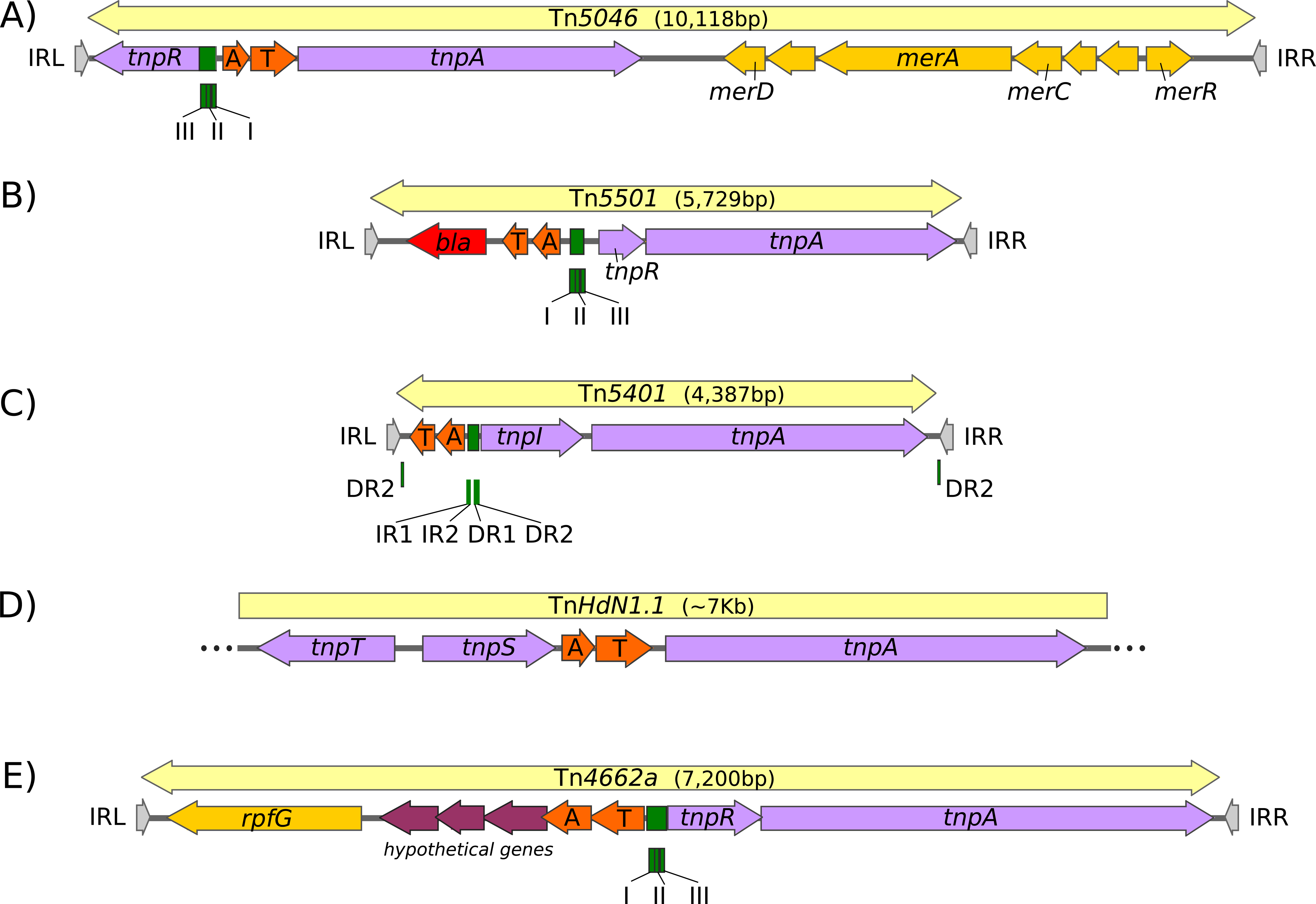
Examples of TA gene pair location in a variety of Tn*3* family transposons. The symbols are the same as those described in Figures 1 and 2 with toxin-antitoxin gene pairs shown in bright orange and genes of unknown function in magenta. A) Tn*5046*, accession # Y18360.1, has an unusual structure with the *mer* passenger genes located downstream from the transposase gene. It carries a typical *tnpR* cognate *res* site. B) Tn*5501.6*, accession # MF487840.1, carries a *bla*NPS-1 passenger gene. It carries a typical *tnpR* cognate *res* site. C) Tn*5401*, accession # U03554.1, There are no known passenger genes apart from the TA gene pair. It carries a typical *tnpI* cognate *irs* site with, in addition a copy of the DR2 TnpI binding site close to each end. D) Tn*HdN1.1*, accession # FP929140.1, is treated as a partial copy since the ends of the transposon have not yet been identified. Consequently, no passenger genes except the TA gene pair have been identified. However, Tn*HdN1.1* carries a typical *tnpT*/*tpnS* resolvase pair and the toxin/antitoxin genes are located between the resolvase and the *tnpA* gene. The *rst* site has not yet been defined, but for other transposons with a TnpS/T/*rst* resolution system, it is located between the divergent *tnpS* and *tnpT* genes (8). E) Tn*4662a*: accession # NC_014124.1. This transposon carries a potential metal dependent phosphohydrolase passenger gene and a *tnpR* cognate *res* site. In this case, in contrast to the vast majority of cases, the toxin gene is located upstream of the antitoxin gene.

### A diversity of TA types

We examined the diversity of the TA modules associated with Tn*3* transposons by comparison of the TA protein sequences with the Pfam database using hmmscan from the HMMER suite (24). Candidates with no Pfam match were searched against the PDB_mmCIF70 database (PDB filtered at 70% sequence identity) using HHsearch, a tool for protein remote homology detection based on profile-to-profile comparison (25). In total, 5 toxin families (RelE/ParE, Gp49, PIN_3, PIN, HEPN) and 6 antitoxin families (ParD, HTH_37, RHH_6, Phd/YefM, AbrB/MazE, MNT) were identified (Tables 1 and S1, Figure 3). All of these toxin families except ParE have been associated with ribonuclease activity, either experimentally or by sequence similarity (26) while ParE inhibits gyrase activity by an unknown molecular mechanism (27). The majority of examples were found in two Tn*3* sub-groups: Tn*3* (2 toxin families; 12 PIN_3 and 2 ParE) and Tn*3000* (3 toxin families; 13 ParE, 5 Gp49 and 1 PIN), while 6 members of 4 different toxin families (ParE, Gp49, PIN_3, PIN and HEPN) were found distributed in the other sub-groups (Tn*Posp1*, Tn*HdN1.1*, and Tn*Tsp1* in sub-group Tn*4651*; Tn*Amu2* and Tn*Sku1* in Tn*5393*; and Tn*Sod9* in Tn*21*). There are 7 different toxin-antitoxin pairs: ParE*-*ParD (14 instances), ParE-PhD (2 instances), PIN_3-RHH_6 (13 instances), Gp49-HTH_37 (7 instances), PIN-Phd (1 instance), PIN-AbrB (1 instance) and HEPN-MNT (1 instance).

In general, the TA genes are arranged with the antitoxin located upstream of the toxin gene. However, TA in reverse order in which the toxin gene precedes that of the antitoxin have been described in the literature (18). Among the 39 TA systems associated with the Tn*3*-family transposons, five members of the Tn*3000* sub-group (Table 1, Figure 3) carried TA systems in which the toxin gene precedes that of the antitoxin. These systems are composed of a Gp49 (PF05973) type toxin (T) of the RelE/ParE superfamily and an HTH_37 (PF13744) type antitoxin (A) of the HigA superfamily. These all have the configuration: <A <T tnpR> tnpA> (where the arrowheads point in the direction of transcription) (18, 28). A similar situation is found in the unrelated Tn*4651* sub-group member, Tn*Posp1_p*. which has the configuration: T> A> tnpR> tnpA>.

Two additional members of the Tn*3* sub-group, Tn*5401* and Tn*Bth4*, both with the configuration tnpI> tnpA>, carried a different TA system: a ParE toxin (Pfam: PF05016) and a ParD antitoxin, which appears to lack the DNA-binding domain. Among the TA gene pairs found in members of the Tn*3000* transposon sub-group, a majority of toxins are of the ParE (Pfam: PF05016) type while the potential antitoxins have no Pfam match. Results from HHpred, indicate that these are antitoxins have an RHH-fold similar to that of the classical ParD antitoxin (PDB: 4Q2U_C). It should be noted that this sub-group of Tn (Tn*5501* and its derivatives) are highly related and differ mainly by the passenger genes they carry. With the exception of Tn*5501.12* (discussed below), all Tn*5501* derivatives have identical or nearly identical toxin and antitoxin protein sequences. They were all identified from the non-redundant NCBI nucleotide database using Tn*5501* as query sequence. Interestingly, a single member of the Tn*4651* sub-group, Tn*Tsp1*, also encodes a nearly identical ParE (identical protein sequence)-ParD (2 aminoacid substitutions) TA pair.

### Acquisition and exchange of TA modules

A relevant question is whether these TA modules (Table 1) were acquired once or multiple times during evolution. This question was addressed by phylogenetic analyses of Tn*3*-associated toxins assigned to the same Pfam group, along with the seed sequences used to build the Pfam hidden markov model (Figure 5). If toxins share a recent common ancestor, there are two possible explanations. In cases where the TA module is found in related transposons (with similar tnpA and/or resolvase genes), it is likely that it was first acquired by a transposon that subsequently diverged. Alternatively, for transposons which are generally not related (different *tnpA* family, different resolvase) but which harbour TA modules that are similar at the DNA level, it is likely that the TA module was acquired by recombination with another transposon. Tn*3* toxins sharing their most common ancestor with non-Tn*3* toxins are likely to have been acquired independently. The different TA modules identified and the Tn*3* family members in which they are found are described below.

**Figure 5.**
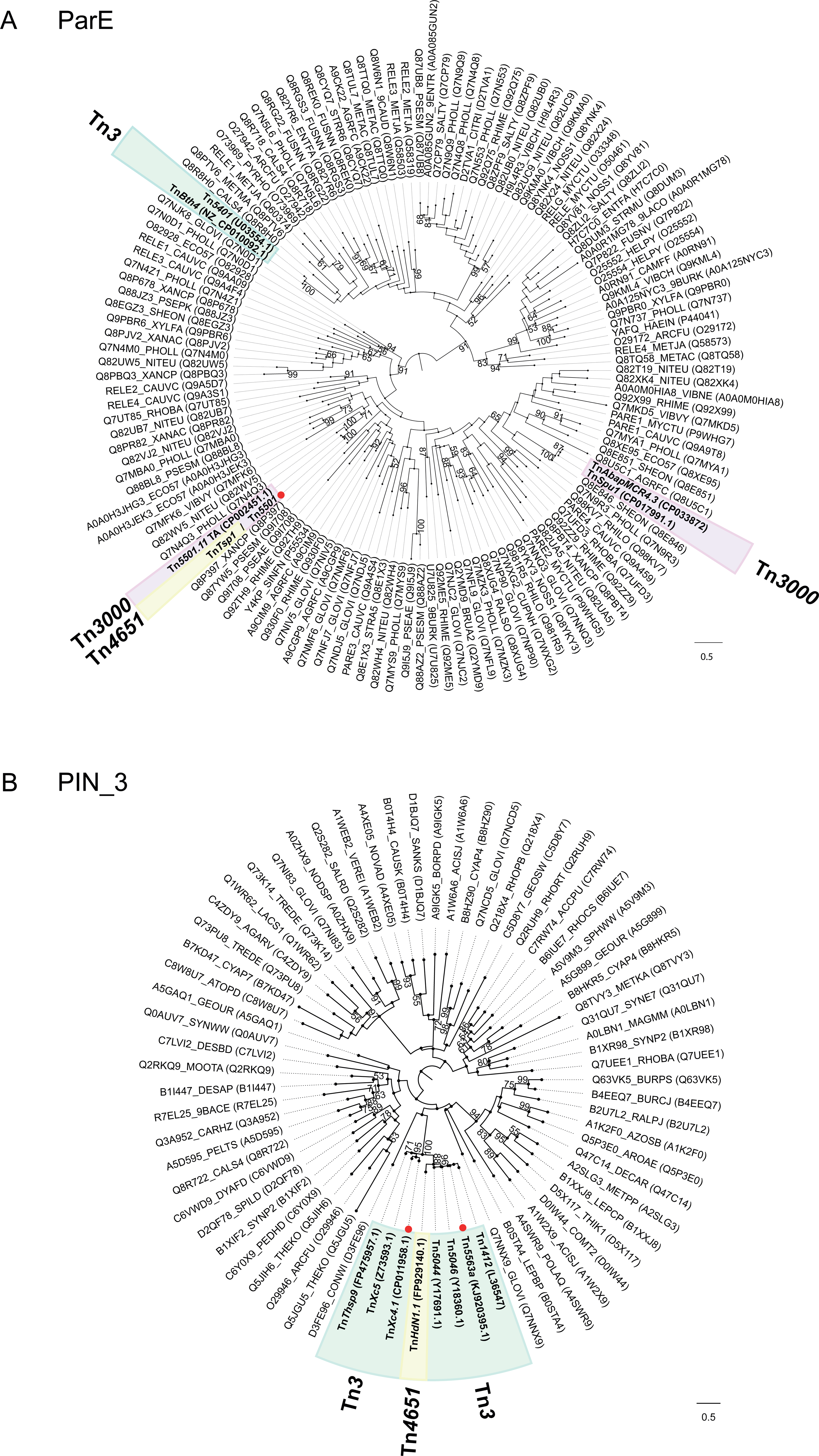
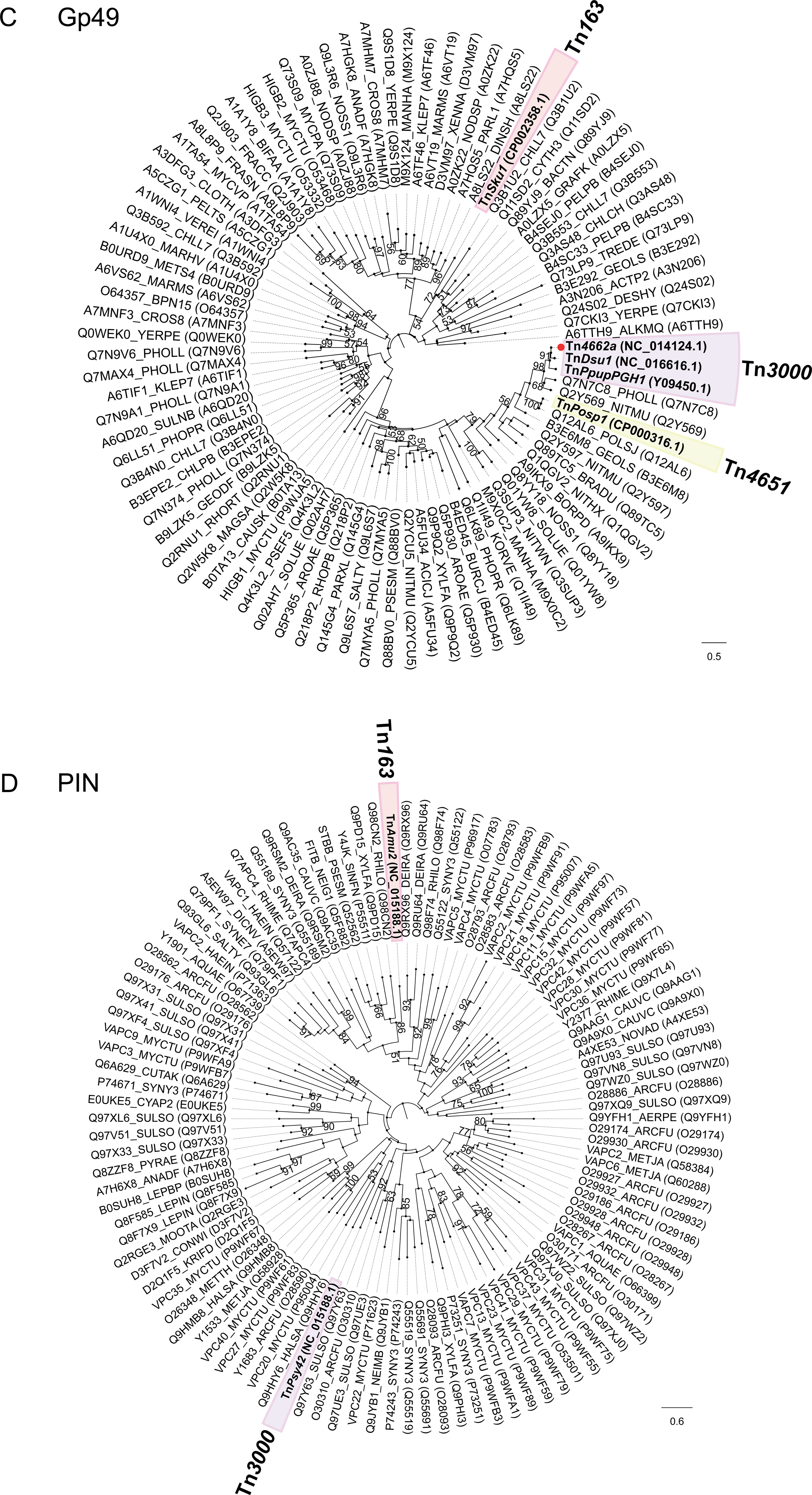
Phylogenetic trees of toxin genes. The phylogenetic history of the transposon-associated toxins reconstructed along the corresponding relative seed proteins downloaded from the Pfam database. The position of transposon-associated toxins is indicated by an outlined colored background indicating the sub-group to which they belong as in Figure 3. Red dots indicate the tips where one toxin sequence was chosen as representative of a set of identical toxins (i.e. there are several Tn examples in the collection). A) ParE. The phylogeny of the ParE toxins suggests that ParE has been recruited 3 times by Tn*3*s: twice by the Tn*3000* sub-group and once by the Tn*3* subgroup. Toxin sequences of Tn*5501*, Tn*5501.1*, Tn*5501.2*, Tn*5501.3*, Tn*5501.4*, Tn*5501.6*, Tn*5501.7*, Tn*5501.8*, *Tn5501.9*, Tn*5501.10* and Tn*Tsp1* are identical, which indicate that the latter acquired the TA module by recombination with a Tn*5501* ancestor. B) PIN_3. The phylogeny of PIN_3 toxins suggests that this gene has been recruited once by the Tn*3* sub-group and further recombined into an ancestor of *TnHdN1.1*. Toxin sequences of Tn*Xc4*, Tn*Xc4.1*, Tn*Xc4.2* and Tn*Xc4.3* are identical. Toxin sequences of Tn*5563a*, Tn*5563a.1* and Tn*5563a.2* are identical. C) Gp49. The phylogeny of Gp49 toxins indicate that these have been recruited in 3 different events by the Tn*3*s: by the Tn*163* sub-group, the Tn*3000* sub-group and the Tn*4651* subgroup. Toxin sequences of Tn*4662a* and Tn*5501.12* are identical. D) PIN. The phylogeny of the PIN toxin suggests that this toxin has been recruited in two separate events.

#### ParE

There are 16 *parE toxin* genes in our collection; fourteen are paired with a *parD* antitoxin gene and two are paired with a *phd/yefM* antitoxin gene (Table 1; Figure 5A). The fourteen *parE*-*parD* modules are found in three transposon sub-families, Tn*3000* (11 examples), Tn*3* (two examples), and Tn*4651* (one example), suggesting that the *parD*-*parE* operon has been acquired three independent times in this collection (Figure 5A).

The first acquisition event concerns the eleven *parE-parD* Tn*3000* sub-group Tns, which are all Tn*5501* relatives and have identical or nearly identical toxin protein sequences (Figure 5A) and significant similarity at the DNA level within the TA modules.

One Tn*5501* derivative, Tn*5501.12*, is an exception since it carries a *gp49-HTH* TA module and is described further below (supplementary figure S1B and D).

The *parE-parD* module located in Tn*Tsp1* (Tn*4651* sub-family) is identical at both the protein and nearly identical at DNA level (95%) with the 10 *parE-parD* modules of the set of Tn*5501* derivatives (Supplementary figure S1D), indicating that the *parD-parE operon* might have been acquired via recombination between a Tn*5501* relative which contributes the DNA segment to the left and an unidentified transposon which contributed the DNA segment to the right to generate Tn*Tsp1*. The recombination point is likely to be at, or close to, the *res* site, where the homology between Tn*Tsp1 res* subsite I and Tn*5501 res* subsite I breaks down (Figure 6A). Resolvase-mediated recombination probably occurs at the dAdT dinucleotide indicated in red in Figure 6A (see (29, 30)).

**Figure 6.**
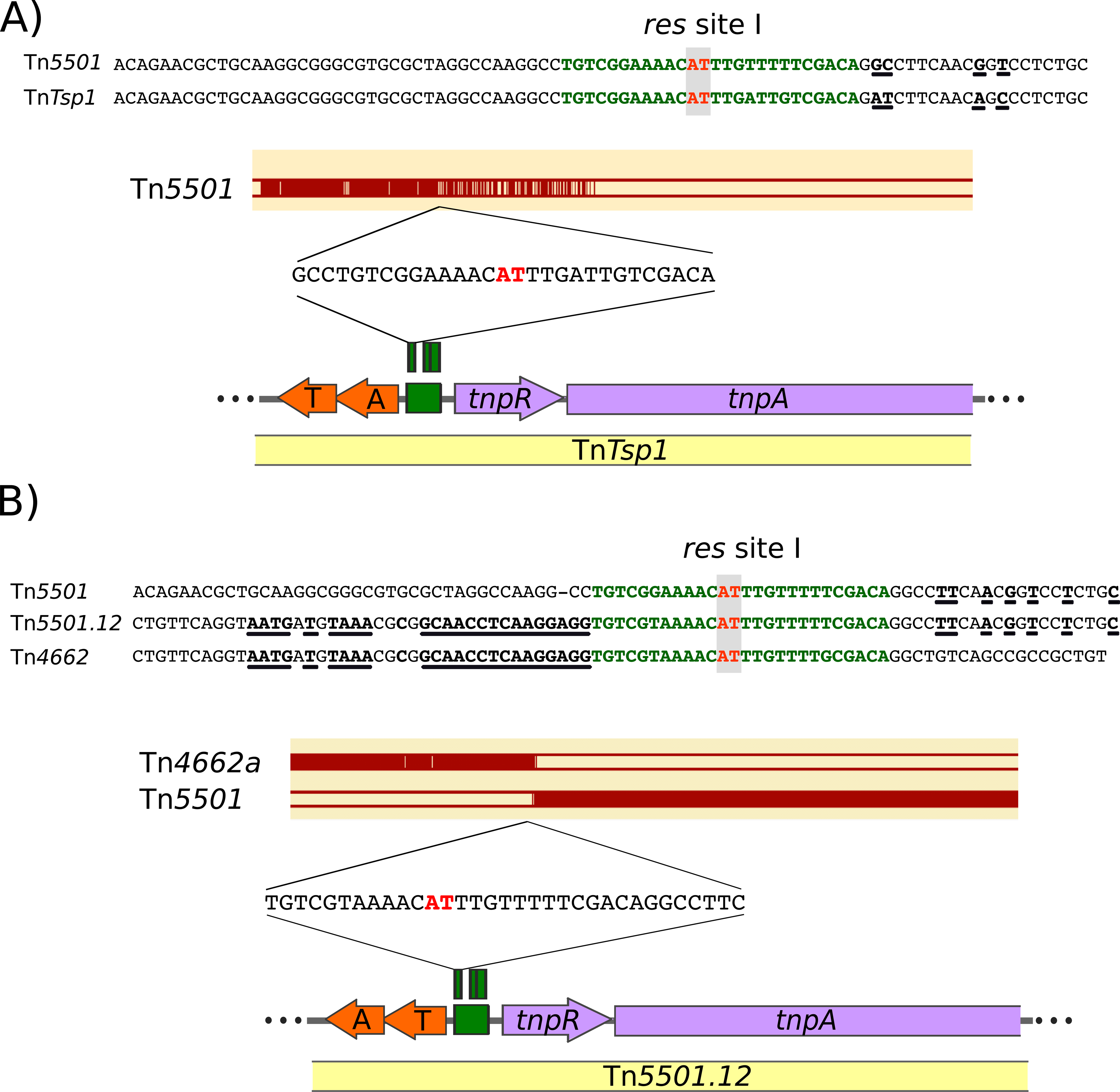
Inter transposon recombination at the res site exchanges TA modules. The symbols are the same as defined in Figures 1, 2 and 4. A) Comparison of Tn*5501* accession # JN648090.1 and Tn*Tsp1* accession # NC_014154 showing a possible recombination point between the two Tn where exchange at the TA gene pair may have occurred. The bottom section shows the region of Tn*Tsp1* including the TA gene module (orange), the *res* site (green), *tnpR* and part of *tnpA* (purple). The top segment shows the equivalent map of Tn*5501*. Below is shown a DNA sequence alignment (magenta) with the equivalent region of Tn*5501*. Both transposons have similar DNA sequences to the left of *res* site I. The level of sequence identity is reduced in *tnpR* and is insignificant in *tnpA*. The *res* site I sequences (green) are shown between the two panels and the AT dinucleotide at which recombination probably occurs is indicated in red. Sequence non-identities are underlined. The two sequences are identical up to the probable recombination site and show some diversity to its right. B) The region of Tn*5501.12* (accession # CP017294.1) showing the 5’ end of the *tnpA* gene, the *tnpR* gene, a *res* site typical of the *tnpR* res sites, and toxin/antitoxin gene pair (note that the toxin gene is upstream of the antitoxin gene (Table 1)). The horizontal magenta lines at the bottom show the alignment of Tn*5501.12* with Tn*5501* (accession # JN648090.1), and Tn*4662a* (NC_014124.1). The right half of Tn*5501* is clearly highly homologous to the right side of Tn*5501.12* whereas the left side of Tn*4662a* is homologous to the left side of Tn*5501.12*. The DNA sequences at the top show the *res* subsite I (green) with the dinucleotide at which recombination should occur in red together with flanking sequences. Underlined bases indicate regions of nucleotide identity. This suggests a scenario in which Tn*5501.12* was generated by recombination at *res* I between transposons similar to Tn*5501* and Tn*4662a*.

The second event is illustrated by Tn*AbapMCR4.3* (31) and Tn*Spu1* (Tn*3000* sub-group) as indicated by the phylogenetic analysis (Figure 5A). Moreover, they are coupled with a different antitoxin, from the Phd/YefM family (Table 1, Figure 3). Identity between the transposition modules () of these two transposons is high intheir transposition modules (including the TA genes) but they differ in their passenger genes.

The third event is represented by two examples, Tn*Bth4* and Tn*5401* (83% identity at the nucleotide level) in the Tn*3* sub-group (Figure 5A). Both carry a TnpI resolvase and they do not have significant similarity with any of the other transposons carrying the *parE*-*parD* TA module (Supplementary Figure S1D).

#### PIN_3

The PIN_3 toxin domain is represented thirteen times (8 unique sequences) in our collection. This domain is associated with an RH_6 antitoxin. Twelve Tns having this module are in the Tn*3* sub-group and one belongs to the Tn*4651* sub-group (Tn*HdN1.1)*. Interestingly, although Tn*HdN*1.1 does not share significant sequence similarity at the nucleotide level with the Tn*3* sub-group Tns featuring the same TA module, at the protein level both toxin and antitoxin from Tn*HdN1.1* share ∼80% sequence identity with those in the Tn*3* sub-group (Figure 5B). Phylogenetic analyses of all thirteen PIN_3 toxins with the seed proteins used to build the corresponding PFAM family suggest that these proteins were recruited by a transposon in a single event and an ancestor of Tn*HdN1.1* acquired the TA module via recombination (Figure 5B).

#### Gp49

Seven TA modules are composed of a Gp49 toxin and an HTH antitoxin. Phylogenetic analyses suggest that this TA module has been recruited in three occasions (Figure 5C).

The first was acquisition by the Tn*3000* sub-group transposons Tn*Dsu1*, Tn*PupPGH1*, Tn*4662a*, Tn*4662a.1* and Tn*5501.12*. Tn*5501.12* is the only relative of Tn*5501* to have this TA module. Sequence comparisons suggest that Tn*5501.12* resulted from exchange of the entire left end of transposon Tn*5501* with a transposon very similar to Tn*4662a* (Figure 6B) carrying a Gp49/HTH_37 TA module. The DNA sequence in this region indicates that recombination between the two transposons occurred at a sequence which resembles *res* site I containing the dinucleotide (in red) at which recombination takes place during the cointegrate resolution step of transposition (29, 30). This mechanism is similar to that proposed for acquisition of the *parE-parD* module by the Tn*4651* sub-group Tn, Tn*Tsp1* as described above. The second acquisition concerns Tn*Posp1* from the Tn*4651* sub-group. The Tn*Posp1* toxin is ∼60 % identical at the protein level to those of the Tn*3000* sub-group are but they are not similar at the DNA level (Supplementary Figure S1B). The phylogenetic reconstruction (Figure 5C) indicates that their most recent common ancestor includes two toxins that are not associated with a Tn*3* transposon, thus suggesting that Tn*Posp1* recruited the TA module independently. The third acquisition event is illustrated by Tn*Sku1* (Tn*163* sub-group), whose toxin does not share a recent ancestor with those of the Tn*3000* group or with the TnPosp1 toxin (Figure 5C).

#### PIN

There are two transposons in the set with a PIN toxin, which appear phylogenetically distant (Figure 5D), suggesting these represent two independent acquisitions. Furthermore, Tn*Amu2* (Tn*4430* sub-group) carries a Phd/YefM antitoxin and Tn*Psy42* has a AbrB/MazE antitoxin.

#### HEPN

Finally, Tn*Sod9* (Tn*21* sub-group), located in the *Shewanella oneidensis* MR-1 megaplasmid, includes an HEPN (<u>H</u>igher <u>E</u>ukaryotes and <u>P</u>rokaryote <u>N</u>ucleotide-binding) type toxin and a MNT (<u>M</u>inimal <u>N</u>ucleotidyl-<u>T</u>ransferase) antitoxin as identified by their similarity to another toxin and antitoxin pair (HHpred hit PDB id 5YEP) encoded in the chromosome of the same strain (32).

Additional indications of independent TA acquisitions are evidenced by Tn*3* derivatives in which the order of genes in the TA module are reversed i.e. the toxin gene being located upstream of the antitoxin gene. This arrangement is found predominantly in members of the Tn*3000* sub-group (Tn*4662a*, Tn*5501.12*, Tn*Dsu1*_p, Tn*Ppup*PGH1, Tn*4662a*.1) although single examples are observed in the Tn*4651* (Tn*Posp1*_p), Tn*21* (Tn*Sod9*) and Tn*163* (Tn*5393.1*) sub-groups.

### The Tn*3*-family-associated TA passenger gene systems are located in a unique position

In most cases, the TA gene pairs are embedded within the transposition module comprising transposase and resolvase genes and the *res* site. They are positioned very close to the *res* sites (see Figure 7). This is in sharp contrast to all other Tn*3* family passenger genes which are generally located away from the resolution and transposon genes and, where known, have often been acquired as integron cassettes or by insertion of other transposons (1). Indeed, several TA-carrying transposons represent closely related derivatives with identical transposase, resolvase and TA modules but contain different sets of passenger genes (e.g. Tn*5501.1* and derivatives .2, .3, .4, …).

**Figure 7.**
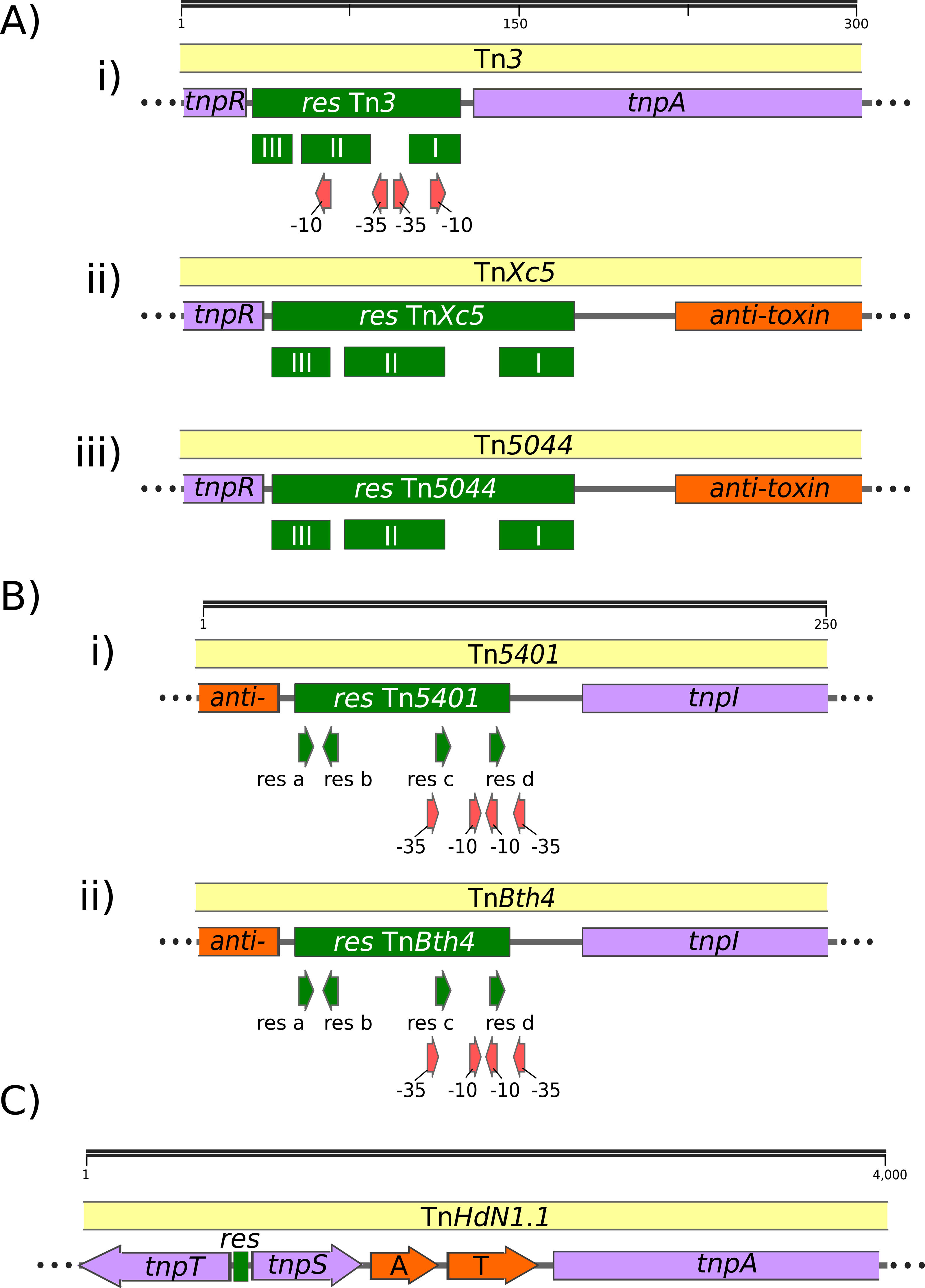
Relationship between the res site, known promoter elements and TA gene pairs. The symbols are identical to those in Figures 1, 2, 4 and 6. In addition, potential or proven minus-10 and minus-35 promoter elements are shown as red arrows. A) *Res* sites (green) with a structure related to Tn*3*. 300 bp including flanking DNA is shown. TnpR (purple) is expressed to the left, and TnpA (purple, Tn*3*) and the toxin/antitoxin genes (orange, Tn*Xc5* and Tn*5044*) to the right. In this type of organization, the res III subsite is proximal to *tnpR*. Recombination leading to cointegrate resolution occurs at a TA dinucleotide within *res* site I.

i. Tn*3* (V00613) *res* site. Taken from Heffron et al. 1979 (12). The *res* site was defined by footprinting using TnpR and by functional deletion analysis. The promoter elements are predicted. ii) Tn*Xc5 res* site (Z73593), also called IS*Xc5*. Taken from Liu et al. (58). The *res* site was defined by footprinting with TnpR. iii) Tn*5044* (Y17691.1) (59–61). The *res* site was defined by comparison with Tn*Xc5* (IS*Xc5*) and as described here. B) Res site organization for transposons carrying res sites for the TnpI resolvase.

i. Tn*5401* (U03554.1) res site. This was identified by footprinting with TnpI and by deletion analysis (5, 12).
ii. Tn*Bth1 res* site (NZ_CP010092.1). Tn*Bth1* is similar but not identical to Tn*5401* over the *res* site but varies considerably in the *tnpI* and *tnpA* genes. It maintains the promoter elements (red arrows) identified in Tn*5401*. *tnpI* and *tnpA* are expressed to the right. The toxin antitoxin pair is expressed to the left. C) Gene organization for transposon Tn*HdN1.1* carrying the TnpS/T resolvase.

In the majority of cases (35 of 39), the Tn*3*-associated TA gene pairs are located directly upstream of the resolvase genes (*tnpR* or *tnpI*) (Figure S2). There are only three exceptions to this. The first is the single example of a derivative with the *TnpS*+*TnpT* resolvase, Tn*HdN1.1* (Figure 3), where the TA genes are located between the resolvase tnpS and transposase genes (Figure 7C). In the second, Tn*Sku1* (not shown), the TA genes are located downstream of and transcribed towards *tnpR*, and in the third, a partial transposon copy, Tn*Amu2*_p (not shown), there is a short ORF of unknown function between the divergently transcribed antitoxin and *tnpR* genes.

### Regulation of TA gene expression

Although it is possible that the TA genes are expressed from their own promoter if present, their position might permit expression from native Tn promoter elements. In Tn*3*, which has been examined in detail, transposase and resolvase gene expression is controlled by promoters found within the *res* site located between the two divergent genes (Figure 7ai). Resolvase binding to these sites autoregulates both *tnpR* and *tnpA* expression (13, 14, 33, 34). The location of the TA genes in proximity to the *res* sites raises the possibility that their expression is also controlled by these promoters.

Few of the *res* sites in the collection of TA-associated Tn*3* family members have been defined either experimentally or by sequence comparison. We therefore attempted to identify potential *res* sites using as a guide the canonical *res*-site organisation schematised in Nicolas et al. (1), a *res* site library (kindly provided by Martin Boocock) and RSAT tools (Regulatory Sequence Analysis Tools; http://rsat.sb-roscoff.fr/) (Materials and Methods). This analysis resulted in identification of 27 potential *res* sites (Table S2). Their organisation is shown in Supplementary Figure S2. For transposons with a TnpR resolvase, it is striking that in every single case, the TA genes are located just downstream from *res* site I, whereas *tnpR* is located next to *res* site III. In transposons with divergent *tnpA* and *tnpR* such as Tn*Xc5* and Tn*5563a* (Figure S2, expanded in Figure 7Aaii and iii), the *tnpA* and *tnpR* genes and *res* sites are organized similarly to those of Tn*3* which does not carry TA (Figure 7Ai), except that *tnpA* is separated from *res* by the intervening TA genes. This organisation is also similar in Tn*3* members in which *tnpA* is downstream of *tnpR* and in the same orientation (e.g., Tn*5501* and Tn*Tsp1*, Supplementary Figure S2).

Promoters have been defined in the *res* (irs) site of the *tnpI*-carrying Tn*5401* (5, 12) and *tnpI* and *tnpA* expression is modulated by TnpI binding to the *res* site (12)(Figure 7Bi). The other *tnpI*-carrying transposon with TA genes, Tn*Bth4* (Figure 7Bii), has an identical *res* site and therefore expression is probably regulated in the same way. Again, the potential promoters are pertinently located for driving expression of the TA module.

Finally, transposon Tn*HdN1.1* (Figure 7C) is the only example in our collection of a *tnpS*+*tnpT* transposon carrying a TA module. The *res* site and relevant promoter elements for the divergently expressed *tnpS+tnpT* have been identified between the *tnpS* and *tnpT* genes in transposon Tn*4651* (8, 9). In Tn*HdN1.1*, the TA gene pair is located to the right of *tnpS*, between *tnpS* and *tnpA* and all three genes are oriented in the same direction. Although the exact regulatory arrangement remains to be determined, it seems possible that the promoters in the *res* site regulate expression of the TA gene pair.

These arrangements raise the possibility that some TA gene expression might occur from a *res* promoter and be subject to control by resolvase binding. On the other hand, if the TA genes do carry their own promoters, then these might regulate downstream transposon genes such as *tnpA*. Further experimental studies are necessary to examine the detailed regulation of toxin, antitoxin and other transposon genes.

## Discussion

As part of our efforts to build a fully annotated transposon database (TnCentral https://tncentral.proteininformationresource.org/), we identified and analysed 190 Tn*3* family transposons (Figure 3 and Table S1) and have observed that 39 of these include type II TA passenger genes from several distinct families (Figure 3, Table 1): 5 toxin families (ParE, Gp49, PIN_3, PIN, HEPN) and 6 antitoxin families (DinJ, HTH_37, RHH_6, Phd/YefM, AbrB/MazE, MNT). Several lines of evidence suggest that there have been multiple independent TA acquisition events, namely: (i) the transposons in our collection feature different families of toxins and antitoxin pairs (Table 1); (ii) in some cases the TA gene order is inverted, and (iii) we observed proteins with no significant sequence similarity within the same toxin/antitoxin family but predicted to shar diverged TA gene pairs. Excluding those cases likely to have arisen from intermolecular Tn e a common fold and therefore could not have diverged from previously recombination (Figure 6), and TnHdN1.1, which also appears to have acquired the TA via recombination, as indicated by the toxin tree (Figure 5B), the most parsimonious interpretation of these observations is that the modules were acquired in 10 separate events. These include three for *parE*, two for PIN3, three for *gp49*, two for PIN and one for HEPN (Table1). At present, it is unclear how such multiple acquisitions have occurred at the molecular level.

In contrast to other passenger genes in Tn*3* family transposons, the TA genes are located at an unusual position within the transposon. They are close to the DNA resolution site (*res*, *irs*, *rst*) (Figure 4) and, more precisely for those with TnpR resolvases, they consistently neighbor *res* site I (Figure S2), a DNA sequence which not only probably includes part of a promoter but is the point at which recombination occurs resulting in cointegrate resolution. For those transposons in which the *tnpR* and *tnpA* genes are divergently orientated (Figures 4a, 6 and S2), the TA module is located between the two genes and expressed in the same direction as *tnpA*. For those in which *tnpR* precedes *tnpA* in the same orientation, the TA module lies upstream from *tnpR* and is oriented in the opposite orientation (e.g. Table 1, Figure 7 and Figure S2). A similar arrangement occurs for the two examples located on *tnpI*-carrying transposons. Only a single example of a *tnpS+tnpT*-carrying transposon with the TA module was identified and here, the TA module is located between the resolvase gene pair and the transposase gene.

This location, close to the key enzymes involved in transposition, suggests that the role of the TA pair might not simply be to provide a general addiction system that stabilizes the host replicon, generally a plasmid, carrying the transposon. It seems possible that they play a more intimate role in stabilising the associated transposon itself. We note, however, that there are two exceptions to this close association of TA genes to the Tn *res* site. For Tn*Sku1*, the TA genes are located downstream of and expressed towards *tnpR*, while in the partial copy, Tn*Amu2*_p, there is a short ORF between the divergently transcribed antitoxin and *tnpR* genes. This does not appear to be related to the 3-component toxin-antitoxin-chaperone (TAC) systems (35).

Interestingly, type II TA expression, like that of *tnpA* and *tnpR*, is tightly regulated at the transcriptional level. Where analysed the toxin-antitoxin complex binds to palindromic sequences located in the operon promoter via the antitoxin DNA-binding domain and acts as a negative transcriptional regulator. This regulation depends critically on the relative levels of toxin and antitoxin in a process known as conditional cooperativity, a common mechanism of transcriptional regulation of prokaryotic type II toxin–antitoxin operons in which, at low toxin/antitoxin ratios, the toxin acts as a co-repressor together with the antitoxin. At higher ratios, the toxin behaves as a de-repressor. It will be important to determine whether the Tn-associated TA genes include their indigenous promotes (18, 36).

In the case of Tn*4631* (19), which is 99% identical to Tn*4662* from plasmid pDK2 (37), the transposon clearly provides a level of stabilisation of the host plasmid. This implies that TA expression occurs in the absence of transposition. There are a number of ways that this could take place (Figure 8). Expression could occur from a resident TA promoter (Figure 8A) if present. However, TA expression might be expected to lead to expression of the downstream *tnpA* gene by readthrough transcription. Alternatively, the absence of a TA promoter, TA expression could occur stochastically from the *res* promoter (Figure 8B). However, this does not rule out the possibility that TA expression is regulated at two levels with a low level “maintenance” expression resulting in the plasmid stabilisation properties described by Loftie-Eaton et al. (19) together with additional expression linked to de-repression of the *tnpA* (and *tnpR*) promoters that must occur during the transposition process (Figure 8C).

**Figure 8.**
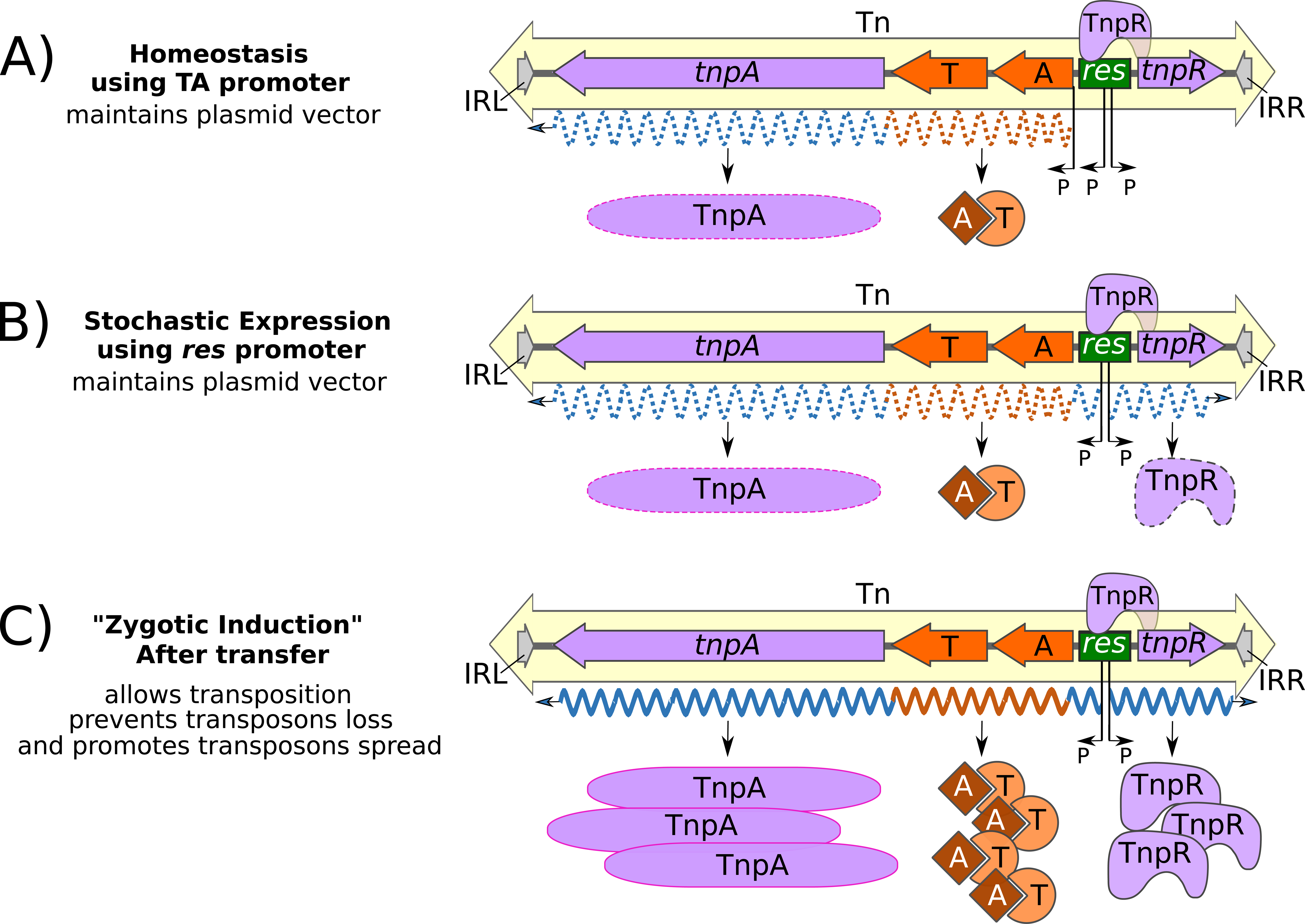
Working Model for the Integration of TA Activity into the Transposition Process. A hypothetical Tn*3* family transposon carrying a TA gene pair is shown. A) Homeostasis on a plasmid stably established in the cell. Transcription (orange and blue dotted wavy line) occurs from a putative endogenous TA promoter (P, proximal to TA) and maintains low toxin (T) and antitoxin (A) levels to maintain the vector plasmid in the host cell population. Expression of *tnpA* and *tnpR* from the *res* site promoters is largely repressed by TnpR binding. However, readthrough transcription from the TA gene pair into tnpA would be expected to result in a level of background TnpA expression. B) If the TA genes do not have an endogenous promoter, stochastic expression (blue and orange dotted wavy lines) from the divergent *res* promoters (P, within the *res* site) would result in low TnpA and TnpR levels as well as low level TA expression. C) Plasmid conjugation into a recipient cell resulting in derepression of the *res* promoters results in higher levels of *tnpA*, *tnpR*, and TA transcription (blue and orange wavy lines) and expression of TA proteins resulting in an increased level of “addiction”.

Indeed, regulation of *tnpR* and *tnpA* by TnpR is a mechanism allowing a burst of TnpA (and TnpR) synthesis, transitorily promoting transposition as the transposon invades a new host. Subsequent repression by newly synthesised TnpR would reduce transposition activity reinstalling homeostasis once the transposon has been established, a process similar to zygotic induction (38) or plasmid transfer derepression as originally observed for ColI (39) and subsequently for R100 (40) and R1 (41). An alternative but non-exclusive explanation stems from the observation that the Tn*6231* TnpR, in addition to the neighboring TA system, enhances plasmid stability (19). Resolvase systems are known to promote resolution of plasmid dimers (see (42)) and it was suggested that integration of the TA system into Tn*6231* “such that all the transposon genes shared a single promoter region” permits coordinated TA and TnpR expression and may facilitate temporary inhibition of cell division while resolving the multimers, promoting plasmid persistence. In this light it is interesting to note that the ccd TA system of the *E. coli* F plasmid is in an operon with a resolvase-encoding gene (43, 44).

Expression of the TA module from the *tnpA*/*tnpR* promoter at the time of the transposition burst could transiently increase invasion efficiency (“addiction”) over and above that provided by the endogenous TA regulation system. If the transposon is on a molecule (e.g., a conjugative plasmid) that is unable to replicate vegetatively in the new host, expression of the TA module without transposition to a stable replicon would lead to loss of the transposon and consequent cell death, whereas cells in which transposition had occurred would survive and give rise to a new population where all cells will contain the Tn. This might be seen as a “take me or die” mechanism, a notion which could be explored experimentally.

Clearly, there remain a number of important questions about the control of TA gene expression that arise from our *in silico* analyses and need to be addressed experimentally. These include whether the TA genes include their own promoters and whether expression is controlled by TA associated promoter elements or by the resident promoters embedded in the *res* sites. Finally, it is an open question whether resolvase binding to *res* represses TA expression either from proximal TA promoters or from *res*-embedded promoters.

## Material and Methods

### Retrieval of prokaryotic genomes and database building

Nucleotide fasta files from complete bacterial and archaeal genomes available in the RefSeq database (45, 46) were downloaded on March 15^th^, 2018. Amino acid sequences of type II toxins and their corresponding antitoxins were retrieved from TADB (47, 48), while Tn*3* transposases and resolvases were retrieved from the ISfinder database (23) and NCBI GenBank (49). These sequences were compiled into multifasta files to be used as databases in subsequent analyses.

### Genomic screening for Tn*3* transposons

The complete genomes were compared to the protein sequences from the TADB, ISfinder and NCBI GenBank databases using tBLASTn 2.2.28 (50) and a custom Python script (Tn*3*finder available from https://tncentral.proteininformationresource.org/TnFinder.html Tn*3*+TA_finder available from https://github.com/danillo-alvarenga/tn3tafinder). ORF prediction was performed with Prodigal 2.6.1 (51) and pre-annotated gbk files were produced with Biopython 1.66 (52). Genome regions presenting translated protein similarity above 40% and 60% of alignment coverage against Tn*3* transposases, resolvases, toxins and antitoxins within maximum distances of 2,000 bp to each other were retrieved for manual curation.

### Manual curation of transposons and accessory genes

Automatic annotations generated by the screening were manually verified in SnapGene Viewer 3.2.1 (https://www.snapgene.com). TA gene pairs were evaluated in greater detail by comparisons with the Pfam 32.0 database (53) using hmmscan from the HMMER 3.1b2 suite (24). Remote homologs were searched with HHpred version 3.2.0 (25) against PDB_mmCIF70, a PDB filtered at 70% sequence identity. HHpred compares the query HMM to the database of HMMs based on PDB chains and generates query-template alignments. Toxin and antitoxin genes with matches against either Pfam or PDB were associated with the name and ID of the Pfam or the PDB ID and known toxins or antitoxins featuring the identified fold (see Table 1).

### Phylogenetic reconstruction

TnpA protein sequences retrieved from our manually curated dataset were aligned with MAFFT 7.309 (54) and their best-fit evolutionary models were predicted with ProTest 3.2.4 (55). A Maximum Likelihood tree was reconstructed with RaxML 8.2.9 (56)) using a bootstrap value of 1,000. The final tree was visualized in FigTree 1.4.4 (http://tree.bio.ed.ac.uk/software/figtree) and edited with Inkscape 0.92.4 (http://www.inkscape.org).

To reconstruct the phylogeny of the toxins, we built a non-redundant toxin set, by removing duplicated sequences. Following the classification of the toxin sequences by comparison with the Pfam database, we downloaded the seed protein sequences for each of the PFAM that matched the toxins. Protein sequence alignment of the toxins with the corresponding PFAM seed sequences and phylogenetic reconstruction followed the same procedure described above for TnpA proteins.

### Sequence comparison between transposons

Transposons were compared all-against-all using blastn. For transposons having a toxin from the same family, all pairwise sequence similarities between the DNA segments comprising the transposase gene, the resolvase and the TA were visualised as dotplots using flexidot version 1.06 (57) with 10 as wordsize (default).

## Acknowledgements

This work was primarily funded by the Global Emerging Infections Surveillance (GEIS) and Response System (P0020_18_WR; awarded to Professor Michael Chandler and Dr. Patrick Mc Gann). We would like to thank Martin Boocock for providing information concerning the *res* sites, Jian Zhang (Protein Information Resource, PIR Georgetown University), Hongzhan Huang (Protein Information Resource, PIR University of Delaware) and Cathy Wu (Protein Information Resource, PIR Georgetown University and University of Delaware) for providing essential expert assistance and both conceptual and practical support. Invaluable contributions were made by Erik Snesrud and Patrick McGann (Walter Reed Army Institute of Research).

## Legends to Supplementary Material

**Figure S1. Pairwise DNA sequence alignment visualisation by dot plot.**

For all transposons with a toxin classified in the same family, the DNA segment that included the transposase gene (large mauve square), the resolvase gene (small mauve square) and the TA module were compared. For visualisation purposes, the size of the TA module was magnified 150-fold. A) Pairwise comparison of sixteen transposons encoding a ParE toxin. The dotplot suggests three independent acquisitions of the TA module, corresponding to: 1) Tn*5501*, its derivatives and Tn*Tsp1*; 2) Tn*Bth4* and Tn*5401*; and 3) Tn*Spu1* and Tn*AbapMCR4.3*. Notice that Tn*Tspu1* is a recombinant since the DNA sequence similarity is limited to the TA module. B) Transposons encoding the PIN_3 toxin. The dot plot suggests that all transposons except Tn*HdN1.1* share sequence similarity over the TA module, thereby suggesting two independent acquisition events of the PIN_3/RHH module. C) Transposons encoding the Gp49 toxin. Seven TA modules are composed of a Gp49 toxin and an HTH antitoxin. The dot plot indicates that this TA module has been recruited on 3 occasions: 1) by the Tn*3000* sub-group transposons Tn*Dsu1*, Tn*PupPGH1*, Tn*4662a*, Tn*4662a.1* and Tn*5501.12*; 2) by Tn*Posp1* (Tn*4651* sub-group) and 3) by Tn*Sku1* (Tn*5393* sub-group). D) The two transposons encoding the PIN toxin do not share any sequence similarity suggesting the TA module was recruited independently on two occasions.

**Figure S2. Different types of transposase, resolvase and TA gene arrangements.**

This figure illustrates the different types of gene arrangement in members of the Tn*3* family which carry toxin/antitoxin genes. Transposons are shown as pale yellow boxes together with transposon names. Recombination sites (*res, irs, rst*) are shown in green, transposition genes in purple and toxin/antitoxin genes in orange. The toxin/antitoxin pair and resolvase types are indicated at the right of each transposon together with the Tn*3* sub-group to which they belong.

**Table S1. Tn*3* family transposons used in this study.**

Column 1: Transposon names and accession numbers; column 2: * transposons with TnpA only and lacking a resolvase gene; column 3: Tn*3* sub-group; column 4: * transposons with TA modules; column 5: the type of resolvase; column 6: bacterial source where known and whether the transposon is located on the chromosome or on a plasmid

**Table S2. Predicted res sites of TA-carrying transposons.**

Column 1: TA-carrying transposon names and accession numbers; column 2: Tn*3* sub-group; columns 3, 5 and 7: nucleotide within res at which res sites I, II and III start (the nucleotide position of the motif, relative to the first nucleotide of the intergenic sequence upstream of *tnpR*); column 4, 6 and 8: DNA sequence of res sites I, II and III; upper case indicates the motif, lower case letters indicate nucleotides that are not part of the motif.

