## Supplementary figures and images for "Toxin-antitoxin gene pairs found in Tn*3* family transposons appear to be an integral part of the transposition module"

### Supplemental figure 1

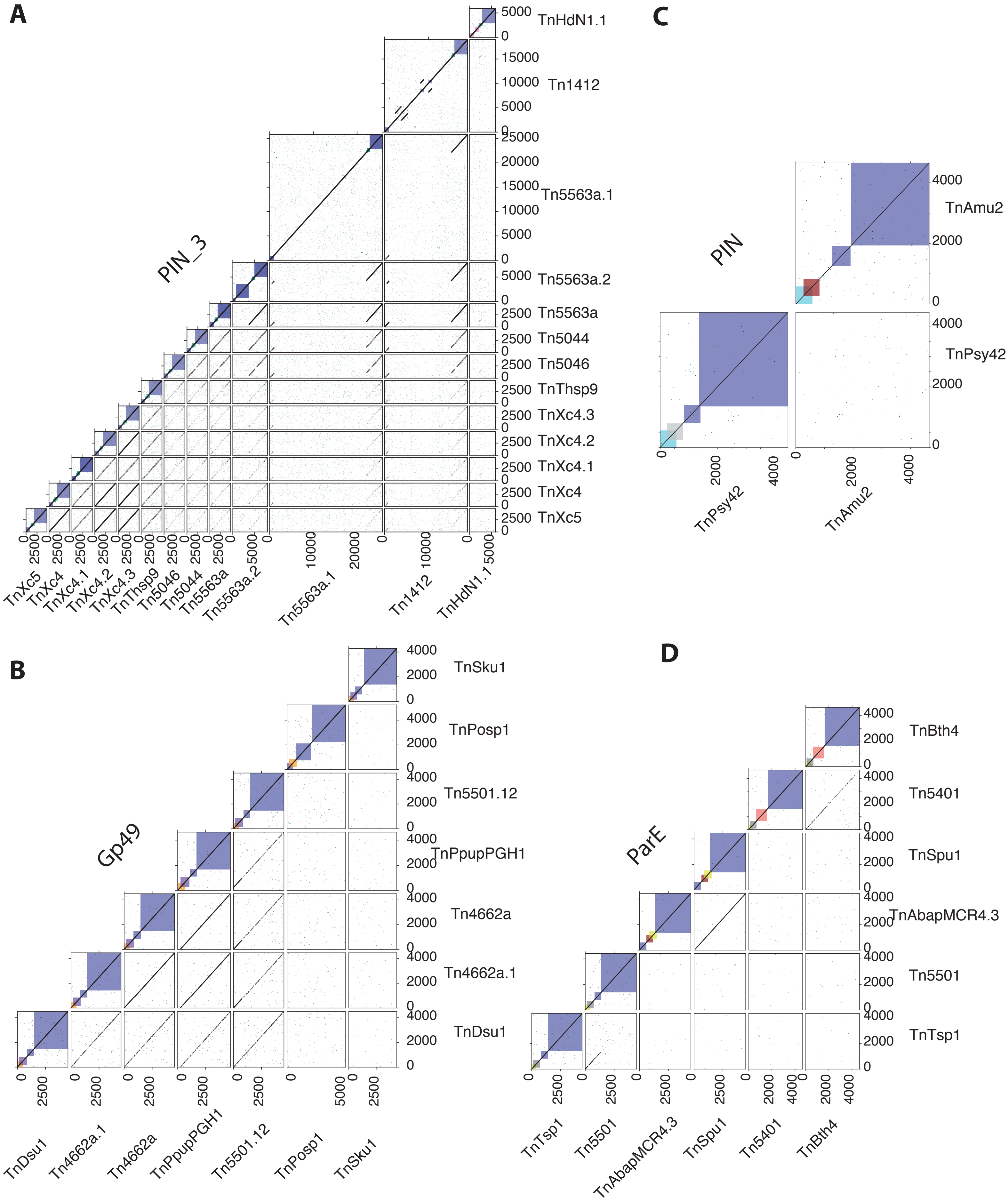

### Supplemental figure 2

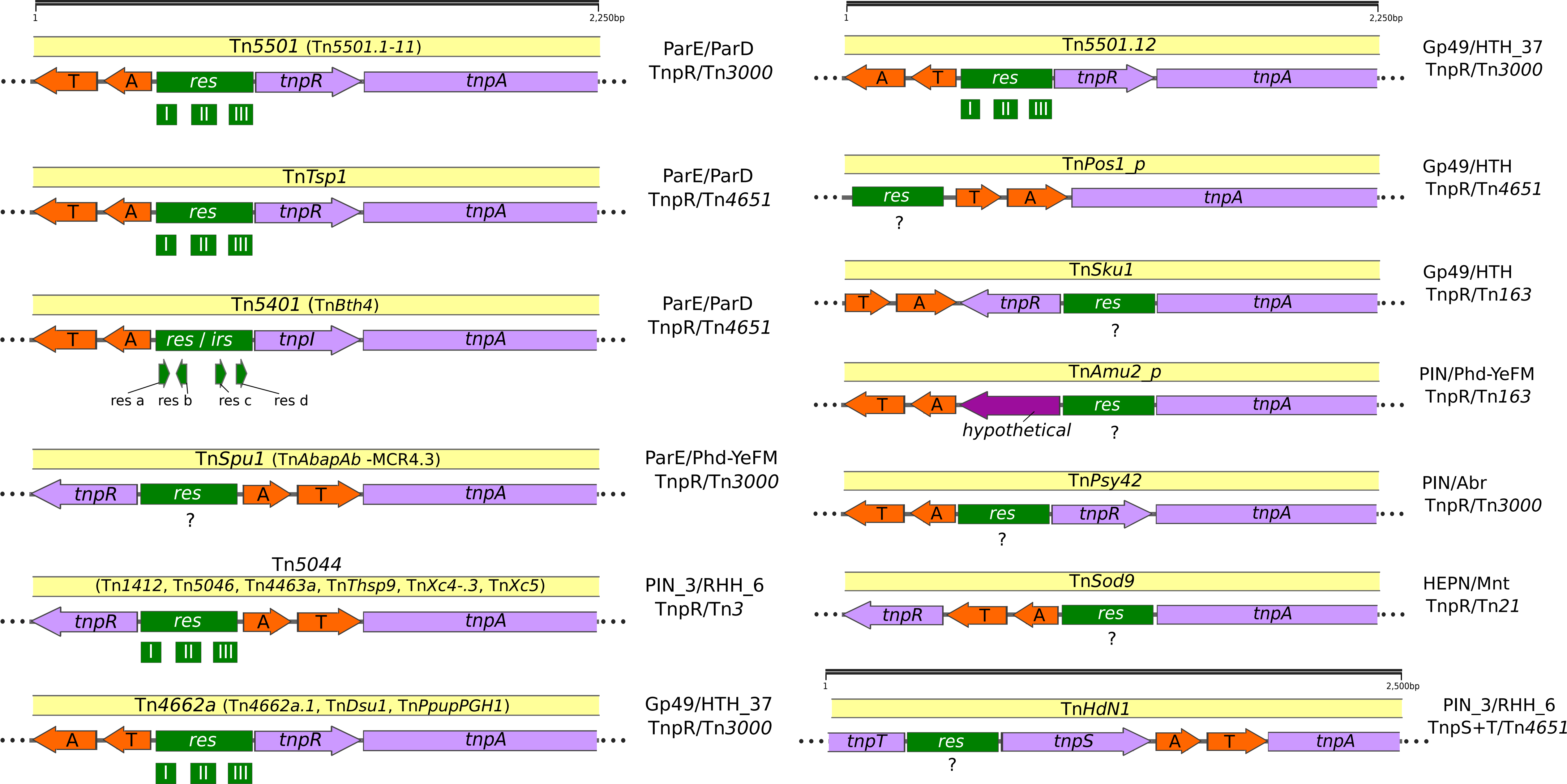
