## Supplemental table S2 for "Toxin-antitoxin gene pairs found in Tn*3* family transposons appear to be an integral part of the transposition module"

**Predicted res sites of TA-carrying transposons**

| **Transposon** | **Sub-Group** | **Start RI** | **RI** | **Start RII** | **RII** | **Start RIII** | **RIII** |
| --- | --- | --- | --- | --- | --- | --- | --- |
| Tn*4662a*_TA_NC_014124.1 | Tn*3000* | 16 | cTGTTCAGGTAATGatGTAAACGCGGCAAC | 53 | aggTGTCGTAAAACAtttGTTTTGCGACAGG | 122 | TCAAAAATGTACGGAAAACTCTATC |
| Tn*5044*_TA_Y17691.1 | Tn*3* | 62 | GTTCGACAAAACGaaGGTTTTGCCGAAC | 123 | ccGCGCTACGGTTCaaataactggtacGCTTAATCAAAGtt | 169 | ATTACTAACGTAGTCGAGTAAACGGTACT |
| Tn*5046*_TA_Y18360.1 | Tn*3* | 62 | GTTCAAAAAAACCttCGTTTTGCCGAAC | 123 | CTGCGCTACGGTTCAAATAACTGGTACGCTTAATCAAAGTT | 169 | ATTACTAACGTAGTCGAGTAAACGGTACT |
| Tn*5393*.1_TA_MF487840.1 | Tn*3000* | 101 | gCCTGTCGGAAAACatTTGTTTTGCGACA | 163 | TGGCCGCAAAATTgtgcggaaaaCTCTGTCGCCAGAcgctacca | 207 | TGCGGAAACCCCTTCTTGATGGTTT |
| Tn*5501*__TA_JN648090.1 | Tn*3000* | 101 | gCCTGTCGGAAAACatTTGTTTTTCGACA | 163 | TGGCCGCAAAATTgtgcggaaaaCTCTGTCGCCAGAcgctacca | 207 | TACGGAAACCTCGTCTTAATGGTTT |
| Tn*5501*.1_TA_CP016447.1 | Tn*3000* | 101 | gCCTGTCGGAAAACatTTGTTTTTCGACA | 163 | TGGCCGCAAAATTgtgcggaaaaCTCTGTCGCCAGAcgctacca | 207 | TACGGAAACCTCGTCTTAATGGTTT |
| Tn*5501*.10_TA_MH053445 | Tn*3000* | 101 | cCCTGTCGGAAAACatTTGTTTTTCGACA | 163 | TGGCCGCAAAATTgtgcggaaaaCTCTGTCGCCAGAcgctacca | 207 | TACGGAAACCTCGTCTTAATGGTTT |
| Tn*5501*.11_TA_CP002451 | Tn*3000* | 101 | gCCTGTCGGAAAACatTTGTTTTTCGACA | 163 | TGGCCGCAAAATTgtgcggaaaaCTCTGTCGCCAGAcgctacca | 207 | TACGGAAACCTCGTCTTAATGGTTT |
| Tn*5501*.12_TA_CP017294.1 | Tn*3000* | 54 | GGTGTCGTAAAACatTTGTTTTTCGACA | 115 | TGGCCGCAAAATTgtgcggaaaaCTCTGTCGCCAGAcgctacca | 159 | TACGGAAACCTCGTCTTAATGGTTT |
| Tn*5501*.2_TA_JN648090.1 | Tn*3000* | 101 | gCCTGTCGGAAAACatTTGTTTTTCGACA | 163 | TGGCCGCAAAATTgtgcggaaaaCTCTGTCGCCAGAcgctacca | 207 | TACGGAAACCTCGTCTTAATGGTTT |
| Tn*5501*.3_TA_GQ983559.1 | Tn*3000* | 101 | gCCTGTCGGAAAACatTTGTTTTTCGACA | 163 | TGGCCGCAAAATTgtgcggaaaaCTCTGTCGCCAGAcgctacca | 207 | TACGGAAACCTCGTCTTAATGGTTT |
| Tn*5501*.4_TA_KY206932.1 | Tn*3000* | 101 | aCCTGTCGGAAAACatTTGTTTTTCGACA | 163 | TGGCCGCAAAATTgtgcggaaaaCTCTGTCGCCAGAcgctacca | 207 | TACGGAAACCTCGTCTTGATGGTTT |
| Tn*5501*.5_A_EF628291.1 | Tn*3000* | 101 | gCCTGTCGGAAAACatTTGTTTTTCGACA | 163 | TGGCCGCAAAATTgtgcggaaaaCTCTGTCGCCAGAcgctacca | 207 | TACGGAAACCTCGTCTTAATGGTTT |
| Tn*5501*.6_TA_MF487840.1 | Tn*3000* | 101 | gCCTGTCGGAAAACatTTGTTTTGCGACA | 163 | TGGCCGCAAAATTgtgcggaaaaCTCTGTCGCCAGAcgctacca | 207 | TGCGGAAACCCCTTCTTGATGGTTT |
| Tn*5501*.7_TA_CP021651.1 | Tn*3000* | 101 | gCCTGTCGGAAAACatTTGTTTTTCGACA | 163 | TGGCCGCAAAATTgtgcggaaaaCTCTGTCGCCAGAcgctacca | 207 | TACGGAAACCTCGTCTTAATGGTTT |
| Tn*5501*.8_TA_KC771559 | Tn*3000* | 101 | gCCTGTCGGAAAACatTTGTTTTTCGACA | 163 | TGGCCGCAAAATTgtgcggaaaaCTCTGTCGCCAGAcgctacca | 207 | TACGGAAACCTCGTCTTAATGGTTT |
| Tn*5501*.9_TA_AJ863570.1 | Tn*3000* | 101 | aCCTGTCGGAAAACatTTGTTTTTCGACA | 163 | TGGCCGCAAAATTgtgcggaaaaCTCTGTCGCCAGAcgctacca | 207 | TACGGAAACCTCGTCTTGATGGTTT |
| Tn*5563a*_TA_KJ920395.1 | Tn*3* | 62 | GTTCGACAAAACGaaGGTTTTGCCGAAC | 123 | ccGCGCTACGGTTCaaataactggtacGCTTAATCAAAGtt | 169 | ATTACTAACGTAGTCGAGTAAACAGTACT |
| Tn*PpupPGH1*_TA_Y09450.1 | Tn*3000* | 42 | cTGTTCAGGTAATGatGTAAACGCGGCAAc | 79 | aggTGTCGTAAAACAtttGTTTTGCGACAGG | 146 | TCAAAAATGTACGGAAAACTCTATC |
| Tn*Thsp9*_TA_FP475957.1 | Tn*3* | 37 | GTTCGATAAAACCttCGTTTTACAAGAC | 100 | CCGCAATTCGTCTaactttgtggtttGATGAATCAAAGtt | 144 | ATTATTCTCTTTCAACGAGTAAAGCAGACT |
| Tn*Tsp1*_TA_NC_014154 | Tn*3000* | 101 | gCCTGTCGGAAAACatTTGATTGTCGACA | 163 | TGGCCGCAAAATTgtgcggaaaaCTGTGTCGCCAGAcgctacca | 207 | TGCGGAAACCGCGTCTTGATGGTTT |
| Tn*Xc4.1*_TA_CP011958.1 | Tn*3* | 67 | GTTCGATAAAACGatCGTTTATCTGAAC | 130 | CAACAAGCAGTCTataaaagtggtttGCATAATCAAAGtt | 174 | ATTGTCTTAATCAACGAGTAAAACGGAAT |
| Tn*Xc4.2*_TA_CP023287 | Tn*3* | 71 | GTTCAATAAAACGatCGTTTTTATGAAC | 134 | CAACAAGCAGTCtgtaatagtggtttgTACAATCAAAGTT | 178 | ATTATTAAAAATTAACGAGTAAAGCAAAAT |
| Tn*Xc4.3*_TA__CP020887.1 | Tn*3* | 70 | GTTCAATAAAACGAatCGTTTTTATGAAC | 133 | CAACAAGCAGTCtgtaatagtggtttgTACAATCAAAGTT | 177 | ATTATTAAAAATTAACGAGTAAAGCAAAAT |
| Tn*Xc4*_TA_Z73593 | Tn*3* | 72 | GTTCAATAAAACGatCGTTTTTATGAAC | 135 | CAACAAGCAGTCtgtaatagtggtttgTACAATCAAAGTT | 179 | ATTATTAAAAATTAACGAGTAAAGCAAAAT |

Column 1: TA-carrying transposon names and accession numbers; column 2: Tn3 sub-group; columns 3, 5 and 7: nucleotide within res at which res sites I, II and III start (the nucleotide position of the motif, relative to the first nucleotide of the intergenic sequence upstream of *tnpR*); column 4, 6 and 8: DNA sequence of res sites I, II and III; upper case indicates the motif, lower case letters indicate nucleotides that are not part of the motif.
