## Supplementary table S1 for "Toxin-antitoxin gene pairs found in Tn*3* family transposons appear to be an integral part of the transposition module"

| **Transposon** | **tnpA only** | **subgroup** | **TA** | **Tnp res** | **Source** |
| --- | --- | --- | --- | --- | --- |
| IS882 NC_005241.1 | * | Tn1071 |  | none | Ralstonia eutropha pHG1 |
| ISBmu19 AP009387.1 |  |  |  |  | Burkholderia multivorans ATCC 17616 |
| ISBmu13 AP009387.1 |  |  |  |  | Burkholderia multivorans ATCC 17616 |
| ISBusp1 NC_007509.1 | * | Tn1071 |  | none | Burkholderia sp. 383 chromosome 3 |
| ISEC63 (FN868832.1) |  | Tn3 |  | R | Escherichia coli p1967-1 |
| ISVsa19 NC_011314.1 | * | Tn3000 |  | none | Aliivibrio salmonicida LFI1238 pVSAL320 |
| Tn1 NC_008357 |  | Tn3 |  | R | Pseudomonas aeruginosa pBS228 |
| Tn1 Mer GQ160960.2 |  | Tn3 |  | R | Serratia marcescens R934 |
| Tn2 KT002541 |  | Tn3 |  | R | Escherichia coli strain HeB7 pHeBE7 |
| Tn2.1-CP028717 |  | Tn3 |  | R | Klebsiella pneumoniae SCM96 pSCM96-1 |
| Tn3 V00613 |  | Tn3 |  | R | Salmonella enterica Wien pZM-3 |
| Tn4 KY749247.1 |  | Tn21 |  | R | Salmonella enterica Paratyphi B pR1 |
| Tn20 AF457211.1 |  | Tn21 |  | R | Escherichia coli |
| Tn21 AF071413 |  | Tn21 |  | R | Escherichia coli plasmid R100/NR1 |
| Tn21.1-MH257753 |  | Tn21 |  | R | Salmonella enterica 1007 pST1007-1A |
| Tn21.2-MH626558 |  | Tn21 |  | R | Salmonella enterica ST1007 pST1007-1B |
| Tn163 L14931 |  | Tn163 |  |  | Rhizobium leguminosarum viciae plasmid |
| Tn501 Z00027 |  | Tn21 |  | R | Pseudomonas aeruginosa pVS1 |
| Tn511 EU287476.1 |  | Tn163 |  | R | Proteus mirabilis R772 |
| Tn551 Y13600.1 |  | Tn4430 |  | R | Staphylococcus aureus pI258 |
| Tn917 M11180 |  | Tn4430 |  | R | Enterococcus faecalis strain DS16 pAD1 |
| Tn1000 X60200.1 |  | Tn3 |  | R | plasmid F |
| Tn1013 AM261760 |  | Tn21 |  | R | pBS228 |
| Tn1071 M65135 |  | Tn1071 |  | R | Comamonas testosteroni strain BR60 |
| Tn1071 composite AB049198.1 | * | Tn1071 |  | R | Delftia acidovorans plasmid pUO1 |
| Tn1331-KC354802.1 |  | Tn3 |  |  | Klebsiella pneumoniae Kp002 pJEG012 |
| Tn1332 DQ174113.1 |  | Tn3 |  | R | Pseudomonas putida |
| Tn1403 AF313472.2 |  | Tn21 |  | R | Pseudomonas aeruginosa pRPL11 |
| Tn1412_TA L36547 |  | Tn3 | * | R | Pseudomonas aeruginosa 2293E |
| Tn1546 M97297.1 |  | Tn4430 |  | R | Enterococcus faecium BM4147 |
| Tn1546_p.1 KR349520.1 |  | Tn4430 |  | R | Enterococcus faecium VREfm 320/07 |
| Tn1546.2 AB247327 |  | Tn4430 |  | R | Enterococcus faecalis plasmid pSL1 |
| Tn1696 U12338.3 |  | Tn21 |  | R | Pseudomonas aeruginosa pR1033 |
| Tn1696.1-CP000451.1 |  |  |  |  | Nitrosomonas eutropha C91 Plasmid1 |
| Tn1721.1 HQ730118.1 |  | Tn21 |  | R | Klebsiella pneumoniae WM2b02 |
| Tn1721 X61367.1 |  | Tn21 |  | R | Gram-negative bacteria |
| Tn2501_p M15197 & Y00502.1 |  | Tn163 |  | R | Escherichia coli K12 |
| Tn2610 AB207867.1 |  | Tn21 |  | R | Escherichia coli pCS200 |
| Tn2815-NC_001275.1 |  |  |  | R | Acetobacter aceti pAC5 |
| Tn3000 AF174129 |  | Tn3000 |  | none | Escherichia coli pMSP071 |
| Tn3434 AY232820.1 |  | Tn163 |  | R | Paracoccus pantotrophus DSM 11072 |
| Tn3926 |  |  |  | R | Yersinia enterolytica |
| Tn4378 CP000355.2 |  | Tn21 |  | R | Cupriavidus metallidurans CH34 pMOL28 |
| Tn4378.1 EU287476 |  |  |  |  | Proteus mirabilis plasmid R772 |
| Tn4380 CP000354.2 |  | Tn21 |  | R | Cupriavidus metallidurans CH34 pMOL30 |
| Tn4401a KT378596.1 |  | Tn3000 |  | R | Klebsiella pneumoniae strain Kp512 plasmid |
| Tn4401b KT378597.1 |  | Tn3000 |  | R | Klebsiella pneumoniae strain Kp15 plasmid |
| Tn4401d-JN974188 |  |  |  | R | Klebsiella pneumoniae plasmid KP529-1 |
| Tn4430 X07651.1 |  | Tn4430 |  | I | Bacillus thuringiensis |
| Tn4556 M29297.1 |  | Tn3000 |  | R? | Streptomyces fradiae |
| Tn4651 |  | Tn4651 | * | R? | Pseudomonas TOL plasmid pWWO |
| Tn4652 AF151431.1 |  | Tn4651 |  | S/T | Pseudomonas putida |
| Tn4652.1 NC_003350 |  | Tn4651 |  | S/T | Pseudomonas putida plasmid pWW0 |
| Tn4653 NC_003350 |  |  |  |  | Pseudomonas putida plasmid pWW0 |
| Tn4656 NC_008275.1 | * | Tn21 |  | R | Pseudomonas putida MT53 plasmid pWW53 |
| Tn4659 NC_008275.1 |  | Tn21 |  | R | Pseudomonas putida MT53 plasmid pWW53 |
| Tn4661 AB375440.1 |  | Tn4651 |  | S/T | Pseudomonas aeruginosa pRms148 |
| Tn4662a.1 TA AY831462.1 |  | Tn3000 | * | R | Pseudomonas putida plasmid pKW1 |
| Tn4662a_TA NC_014124.1 |  | Tn3000 | * | R | Pseudomonas putida plasmid pDK1 |
| Tn5036 Y09025 |  | Tn21 |  | R | Enterobacter cloacae |
| Tn5037 AJ251743.1 |  | Tn21 |  | R | Thiobacillus ferrooxidans |
| Tn5041 X98999.3 |  | Tn4651 |  | S/T | Pseudomonas sp. |
| Tn5044_TA Y17691.1 |  | Tn3 | * | R | Xanthomonas campestris |
| Tn5045 FN821089.1 |  | Tn21 |  | R | Pseudomonas sp. Tik3 |
| Tn5045.1 NC_008357.1 |  | Tn21 |  | R | Pseudomonas aeruginosa pBS228 |
| Tn5046 TA Y18360.1 |  | Tn3 | * | R | Pseudomonas sp. strain LS46-6 |
| Tn5046.1_TA-CP029092 |  |  |  |  | Pseudomonas aeruginosa AR441 plasmid unnamed2 |
| Tn5060 AJ551280.1 |  | Tn21 |  | R | Pseudomonas sp. A19-1 |
| Tn5070 Y17830.1 |  | Tn3 |  | R | Pseudomonas sp. BW13 |
| Tn5084 AB066362.1 |  | Tn4430 |  | R | Bacillus cereus RC607 |
| Tn5393c AY342395.1 |  | Tn163 |  | R | Pseudomonas syringae syringae pPSR1 |
| Tn5393 M95402.1 |  | Tn163 |  | R | Erwinia amylovora pEa34 |
| Tn5393.1_TA MF487840.1 |  | Tn163 | * | R | Pseudomonas aeruginosa PA34 |
| Tn5393.2 CP030921.1 |  | Tn163 |  | R | Escherichia coli KL53 pKL53-M |
| Tn5393.3 LT985287.1 |  | Tn163 |  | R | Escherichia coli RPC3 plasmid RCS69_pI |
| Tn5393.4 AJ627643.4 |  | Tn163 |  | R | Alcaligenes faecalis |
| Tn5393.5 CP027411.1 |  | Tn163 |  | R | Salmonella enterica FDAARGOS_319 p unnamed |
| Tn5393.6 CU928144.1 |  | Tn163 |  | R | Escherichia fergusonii ATCC 35469 pEFER |
| Tn5393.7 LT827129.1 |  | Tn163 |  | R | Escherichia coli K12 J53 R6K. |
| Tn5393.8 CP002090.1 |  | Tn163 |  | R | Salmonella enterica Kentucky pCS0010A |
| Tn5393.9 KU987453.1 |  | Tn163 |  | R | Klebsiella pneumoniae 05K0261 pF5111 |
| Tn5393.10 CP019905.1 |  | Tn163 |  | R | Escherichia coli MDR_56 p unnamed6 |
| Tn5393.11 CP000602.1 |  | Tn163 |  | R | Yersinia ruckeri YR71 plasmid pYR1 |
| Tn5393e GU562437.2 |  | Tn163 |  | R | Salmonella enterica Typhimurium pSRC125 |
| Tn5401_TA U03554.1 |  | Tn3 | * | I | Bacillus thuringiensis serovar morrisoni EG2158 |
| Tn5403 X75779.1 |  | Tn163 |  | R | Klebsiella pneumoniae (ozonae) |
| 5403a EU278476 |  |  |  |  |  |
| Tn5501 _TA JN648090.1 |  | Tn3000 | * | R | Delftia sp. KV29 plasmid pKV29 |
| Tn5501.1 CP016447.1 |  | Tn3000 | * | R | Acidovorax sp. RAC01 chromosome |
| Tn5501.3 _TA GQ983559.1 |  | Tn3000 | * | R | Uncultured bacterium pAKD4 |
| Tn5501.4_TA KY206932.1 |  | Tn3000 | * | R | Uncultured bacterium clone contig003. |
| Tn5501.5_A EF628291.1 |  | Tn3000 | * | R | Uncultured bacterium plasmid pGNB1 |
| Tn5501.6_TA MF487840.1 |  | Tn3000 | * | R | Pseudomonas aeruginosa PA34 |
| Tn5501.7_TA CP021651.1 |  | Tn3000 | * | R | Acidovorax sp. T1 plasmid p3-T1 |
| Tn5501.8 _TA KC771559 |  | Tn3000 | * | R | Comamonas sp. 7D-2 plasmid pBHB |
| Tn5501.9_TA AJ863570.1 |  | Tn3000 | * | R | Uncultured bacterium IncP-1beta plasmid pB8 |
| Tn5501.10_TA MH053445 |  | Tn3000 | * | R | Pseudomonas aeruginosa PA1280 plasmid pICP-4GES |
| Tn5501.11_TA CP002451 |  | Tn3000 | * | R | Alicycliphilus denitrificans BC plasmid pALIDE02 |
| Tn5501.12_TA CP017294.1 |  | Tn3000 |  | R | Pseudomonas aeruginosa PA83 plasmid unnamed1 |
| Tn5501.13_TA-EF628291 |  | Tn3000 |  | R | Uncultured bacterium plasmid pGNB1 |
| Tn5504 AY365053 |  | Tn3000 |  | R | Ralstonia eutropha JMP134 plasmid pJP4 |
| Tn5563a_TA KJ920395.1 |  | Tn3 | * | R | Pseudomonas mendocina LM7 |
| Tn5563a.1_TA |  | Tn3 | * | R | Pseudomonas aeruginosa MRSN12280 chromosome |
| Tn5563a.2_TA CP028567 |  | Tn3 | * | R | Aeromonas hydrophila plasmid pMCR5_045096 |
| Tn5719-AJ431260 |  | Tn21 |  | R | Uncultured bacterium plasmid pB4 |
| Tn5720-AJ431260 |  | Tn21 | * | R | Uncultured bacterium plasmid pB4 |
| Tn6001_p EF138817.1 |  | Tn21 |  | R | Pseudomonas aeruginosa strain 450 |
| Tn6005 EU591509.1 |  | Tn21 |  | R | Enterobacter cloacae strain JKB7 |
| Tn6025-GU562437 |  | Tn163 |  |  | Salmonella enterica Typhimurium pSRC125 |
| Tn6060 GQ161847 |  | Tn21 |  | R | Pseudomonas aeruginosa 37308 genomic island |
| Tn6061_p GQ388247.1 |  | Tn21 |  | R | Pseudomonas aeruginosa BM4530 |
| Tn6122 JN127372.1 |  | Tn163 |  | R | Paracoccus halophilus strain JCM |
| Tn6134 AB610645.1 |  | Tn163 |  | R | Sphingobium japonicum UT26 |
| Tn6135 AB610646.1 |  | Tn163 |  | R | Sphingobium japonicum UT26 |
| Tn6136 AB610647.1 |  | Tn163 |  | R | Sphingobium japonicum UT26 |
| Tn6137 AB610648.1 |  | Tn163 |  | R | Sphingobium japonicum UT26 |
| Tn6138 AB610649.1 |  | Tn163 |  | R | Sphingobium japonicum UT26 |
| Tn 6231 TA |  |  |  |  |  |
| Tn6238 KJ511462.1 |  | Tn3 |  | R | Klebsiella pneumoniae KpF7 pMPQDC1 |
| Tn6294 LC015492.1 |  | Tn4430 |  | R | Paenibacillus sp. EOA1 |
| Tn6332 LC155216.1 |  | Tn4430 |  | R | Bacillus sp. TW6 |
| TnAau4 NC_008711 | * | Tn3 |  | none | Paenarthrobacter aurescens TC1 |
| TnAbapMCR4.3 CP033872 |  |  |  | R | Acinetobacter baumannii MRSN15313 pAb-MCR4.3 |
| TnAcsp1 MF399199.1 |  | Tn21 |  | R | Acinetobacter baumannii D46 plasmid pD46-4 |
| TnAfe8687-AAS45120.1 | * | Tn21 |  | none | Alcaligenes faecalis NCIB 8687 |
| TnAli20 FQ311873.1 |  | Tn163 |  | R | Azospirillum lipoferum 4B plasmid AZO_p5 |
| TnAmu1 AP012041.1 |  | Tn163 |  |  | Acidiphilium multivorum AIU301 plasmid pACMV6 |
| TnAmu2_p TA NC_015188.1 |  | Tn163 | *? | R | Acidiphilium multivorum AIU301 pACMV4 |
| TnAnox1-KB205935.1 |  | Tn4430 |  | R | Anoxybacillus flavithermus TNO-09.006 |
| TnAO22 EU696790.1 |  | Tn21 |  | R | Achromobacter sp. AO22 |
| TnARS1_p AY780525.1 |  | Tn4430 |  | R | Bacillus sp. MB24 |
| TnArsp6 NC_008538 |  | Tn163 |  | R | Arthrobacter sp. FB24 plasmid 2 |
| TnAs1 CP022426.1 |  | Tn21 |  | R | Aeromonas salmonicida subsp. pectinolytica 34mel |
| TnAs2 JN106175.1 |  | Tn21 |  | R | Uncultured bacterium pAKD34 |
| TnAs3 CP000645.1 |  | Tn21 |  | R | Aeromonas salmonicida A449 plasmid 4 |
| TnAsz17 AP010946.1 |  | Tn163 |  | R | Azospirillum sp. B510 |
| TnAtcArs AY821803.1 |  | Tn21 |  | R | Acidithiobacillus caldus |
| TnAypMCR5-CP028567 |  | Tn21 |  | R | Aeromonas hydrophila WCHAH045096 pMCR5_045096 |
| TnAzs29 NC_013855 |  | Tn163 |  | R | Azospirillum sp. B510 pAB510a |
| TnBmu14 |  |  |  |  |  |
| TnBth1_p extended CP013280 |  | Tn3 |  |  | Bacillus thuringiensis israelensis AM65-52 pAM65-52-5-100K |
| TnBth2 LXLI01000054 |  | Tn4651 |  | R | Bacillus thuringiensis GOE4 BTGOE4_contig000054 |
| TnBth3 CP003766 |  | Tn163 |  | R | Bacillus thuringiensis HD-789 pBTHD789-3 |
| TnBth4 TA NZ_CP010092.1 |  | Tn3 | * | I | Bacillus thuringiensis galleriae HD-29 pBMB126 |
| TnCfrpOZ172-CP016763.1 |  |  |  | R | Citrobacter freundii B38 pOZ172 |
| TnChe1 CP000391.1 |  | Tn163 |  | R | Chelativorans sp. BNC1 plasmid 2 |
| TnDra1 CP015084 ???? |  | Tn4651 |  | R | Deinococcus radiodurans R1 pCP1 |
| TnDsu1_p TA NC_016616.1 |  | Tn3000 | * | R | Dechlorosoma suillum PS |
| TnEc1 KX555451.1 |  | Tn163 |  | R | Escherichia coli pMCR-11EC-P293 |
| TnEcO26-BDIH01000107.1 |  | Tn21 |  | R | Escherichia coli O26 |
| TnHad2 AB049198.1 |  | Tn1071 |  |  | Delftia acidovorans plasmid pUO1 |
| TnHdN1 TA FP929140.1 |  | Tn4651 | * | S/T | gamma proteobacterium HdN1 |
| TnKox2 CP003684 | * | Tn3000 |  | none | Klebsiella michiganensis E718 pKOX_R1 |
| TnLfArs DQ057986.1 |  | Tn21 |  | R | Leptospirillum ferriphilum |
| TnMERI1_p LC152290.1 |  | Tn4430 |  | R | Bacillus megaterium MB1 |
| TnMex22 CP001511.1 |  | none |  | R | Methylobacterium extorquens AM1 megaplasmid |
| TnMex38 CP001513.1 |  | Tn163 |  | R | Methylobacterium extorquens AM1 p2META1 |
| TnMpo10 NC_010721 |  | Tn163 |  | R | Methylobacterium populi BJ001 pMPOP02 |
| TnNpu13 CP001041.1 |  | Tn4651 |  | R | Nostoc punctiforme PCC 73102 pNPUN04 |
| TnOtChr EF469735.1 |  | Tn21 |  | R | Ochrobactrum tritici |
| TnPa38 CP003149.1 |  | Tn21 |  | R | Pseudomonas aeruginosa DK2 |
| TnPa40 CP003149.1 |  | Tn21 |  | R | Pseudomonas aeruginosa DK2 |
| TnPa40.1 CP020704.1 |  | Tn21 |  | R | Pseudomonas aeruginosa strain PASGNDM699 |
| TnPa42 CP003149.1 |  | Tn4651 |  | S/T | Pseudomonas aeruginosa DK2 |
| TnPa43 ISfinder |  | Tn4651 |  | R |  |
| TnpEc158- KY887596.1 |  |  |  |  | Escherichia coli Ec158 pEc158 |
| TnPersp1 CP012854.1 |  | Tn4651 |  | R | Persicobacter sp. JZB09 plasmid JZB09-Plasmid11 |
| TnPosp1_p TA CP000316.1 |  | Tn4651 | *? | R | Polaromonas sp. JS666 |
| TnPpa1 (composite) DQ149577.1 |  | Tn163 |  |  | Paracoccus pantotrophus |
| TnPpupPGH1 TA Y09450.1 |  | Tn3000 | * | R | Pseudomonas putida plasmid pPGH1 |
| TnPsy30 FR820585 |  | Tn3000 |  | R | Pseudomonas savastanoi pv. savastanoi pPsv48A |
| TnPsy39 CP018203.1 |  | Tn3000 |  | R | Pseudomonas syringae actinidiae ICMP 9853 p9853_A |
| TnPsy42 TA KX009060.1 |  | Tn3000 | * | R | Pseudomonas syringae pv. actinidiae RT594 pUR_RT594 |
| TnRsp12 FO082821 |  | none |  | R | Rhizobium sp. NT-26 plasmid NT26_p1 |
| TnSba14 NC_009052 |  | Tn163 |  | R | Shewanella baltica OS155 |
| TnSen1.1 KY807921 |  | Tn21 |  | R | Salmonella enterica enterica Paratyphi B pSE13-SA01718 |
| TnSen1.2 CP028162 |  | Tn21 |  | R | Pseudomonas aeruginosa MRSN12280 |
| TnSF1 AF188331.1 |  | Tn4651 |  |  | Shigella flexneri |
| TnSgr AB605439.1 |  | Tn4651 |  | R | Streptomyces griseus |
| TnShes11 NC_008573 |  | Tn21 |  | R | Shewanella sp. ANA-3 plasmid 1 |
| TnShfr1 NC_008345 |  | Tn21 |  | R | Shewanella frigidimarina NCIMB 400 |
| TnShfr9 NC_008345 | * | Tn3000 |  | none | Shewanella frigidimarina NCIMB 400 |
| TnSku1 TA CP002358.1 |  | Tn163 | * | R | Sulfuricurvum kujiense DSM16994 plasmid pSULKU03 |
| TnSmeSM11a-1 CP021217.1 |  | Tn4651 |  | R | Sinorhizobium meliloti RU11/001 plasmid pSymA |
| TnSod9 TA NC_004349 |  | Tn21 | * | R | Shewanella oneidensis MR-1 plasmid megaplasmid |
| TnSpu1 TA CP017991.1 |  | Tn3000 | * | R | Enterobacter cloacae sp. ECNIH7 pENT-1ac |
| TnStma1_p NC_017671 |  | Tn4651 |  | R | Stenotrophomonas maltophilia D457 |
| TnTF5 NC_005023.1 |  | Tn4651 |  | R | Acidithiobacillus ferridurans ATCC 33020 pTF5 |
| TnThsp9 TA FP475957.1 |  | Tn3 | * | R | Thiomonas sp. str. 3As plasmid pTHI |
| TnTin1 (inactive) CP002021 |  | Tn4651 |  | T only | Thiomonas intermedia K12 pTINT01 |
| TnTsp1 TA NC_014154 |  | Tn4651 | * | R | Thiomonas intermedia K12 plasmid pTINT01 |
| TnXax1 AE008925 |  | Tn4651 |  | S/T | Xanthomonas axonopodis pv. citri str. 306 pXAC64 |
| TnXax1-like AY389509.1 |  | Tn4651 |  | S/T | Xanthomonas euvesicatoria 75-3 |
| TnXax1.1 NC_016053 |  | Tn4651 |  | ? | Xanthomonas arboricola pruni CFBP 5530 pXap41 |
| TnXax1.2 CP002914 |  | Tn4651 |  | S/T | Xanthomonas axonopodis pv. citrumelo F1 |
| TnXc5 TA Z73593 |  | Tn3 | * | R | Xanthomonas campestris |
| TnXc4 TA CP009039.1 |  | Tn3 |  |  | Xanthomonas citri pXCAW58 |
| TnXc4.1 TA CP011958.1 |  | Tn3 | * | R | Xanthomonas oryzae oryzicola CFBP7331 |
| TnXc4.2 TA CP023287 |  | Tn3 | * | R | Xanthomonas citri pv. citri 03-1638-1-1 pP2 |
| TnXc4.3 TA CP020887.1 |  | Tn3 | * | R | Xanthomonas citri TX160149 plasmid unnamed2 |
| TnXca1 NC_007507 |  | Tn4651 | ?? | S/T | Xanthomonas campestris vesicatoria pXCV183, |
| TnXO1 NC_007322.2 |  | Tn4651 |  | R | Bacillus anthracis 'Ames Ancestor' plasmid pXO1 |
| TnXo19 KR071788 |  | Tn4651 |  | S/T | Xanthomonas oryzae pv. oryzicola GX01 pXOCgx01 |
| TnYps3 FM178282.1 |  | Tn4651 |  | R | Yersinia pseudotuberculosis pGDT4 |
| TnDspM7H15-1_p_TA -CP035300.1 |  |  |  |  | Dyella sp. M7H15-1 chromosome |
| TnXtaPLS235_P_TA-CP011800.1 |  |  |  |  | Xylella taiwanensis PLS235 chromosome |

Column 1: Transposon names and accession numbers; column 2: * indicates transposons with TnpA only and lacking a resolvase gene; column 3: Tn3 sub-group; column 4: * indicates transposons with TA modules; column 5: indicates the type of resolvase; column 6: lists the bacterial source where known and whether the transposon is located on the chromosome or on a plasmid
